# Ketogenic Diet Prevents Heart Failure from Defective Mitochondrial Pyruvate Metabolism

**DOI:** 10.1101/2020.02.21.959635

**Authors:** Kyle S. McCommis, Attila Kovacs, Carla J. Weinheimer, Trevor M. Shew, Timothy R. Koves, Olga R. Ilkayeva, Dakota R. Kamm, Kelly D. Pyles, M. Todd King, Richard L. Veech, Brian J. DeBosch, Deborah M. Muoio, Richard W. Gross, Brian N. Finck

## Abstract

The myocardium is metabolically flexible and can use fatty acids, glucose, lactate/pyruvate, ketones, or amino acids to fuel mechanical work. However, impaired metabolic flexibility is associated with cardiac dysfunction in conditions including diabetes and heart failure. The mitochondrial pyruvate carrier (MPC) is required for pyruvate metabolism and is composed of a hetero-oligomer of two proteins known as MPC1 and MPC2. Interestingly, MPC1 and MPC2 expression is downregulated in failing human hearts and in a mouse model of heart failure. Mice with cardiac-specific deletion of MPC2 (CS-MPC2-/-) exhibited loss of both MPC2 and MPC1 proteins and reduced pyruvate-stimulated mitochondrial respiration. CS-MPC2-/- mice exhibited normal cardiac size and function at 6-weeks old, but progressively developed cardiac dilation and contractile dysfunction thereafter. Feeding CS-MPC2-/- mice a ketogenic diet (KD) completely prevented or reversed the cardiac remodeling and dysfunction. Other diets with higher fat content and enough carbohydrate to limit ketosis also improved heart failure in CS-MPC2-/- mice, but direct ketone body provisioning provided only minor improvements in cardiac remodeling. Finally, KD was also able to prevent further remodeling in an ischemic, pressure-overload mouse model of heart failure. In conclusion, loss of mitochondrial pyruvate utilization leads to dilated cardiomyopathy that can be corrected by a ketogenic diet.

---

The myocardium requires vast amounts of chemical energy stored in nutrients to fuel cardiac contraction. To maintain this high metabolic capacity, the heart is extremely flexible and can adapt to altered metabolic fuel supplies during diverse developmental, nutritional, or physiologic conditions. Cardiac mitochondria are capable of oxidizing fatty acids, pyruvate (derived from either glucose or lactate), ketone bodies, or amino acids when needed. Whereas fatty acids are considered a predominant fuel source for normal adult hearts^1, 2^, several physiological conditions can increase the importance of other substrates for cardiac metabolism. For example, the mammalian fetal heart relies mostly on anaerobic glycolysis until oxygen is abundant and the oxidative capacity of the heart matures postnatally^3^. Exercise greatly enhances myocardial lactate extraction and metabolism^4^. Fasting enhances ketone body delivery to the heart, and myocardial ketone extraction and metabolism can be increased in proportion to delivery^5–7^.

A hallmark of heart failure in mice and in humans is a metabolic switch away from mitochondrial oxidative metabolism^8–11^. Fatty acid oxidation (FAO) is reduced in the failing heart as a result of deactivating the expression of a wide transcriptional program for FAO enzymes and transporters^8, 12–14^ and other mitochondrial metabolic enzymes^8, 10, 11^. The deactivation of mitochondrial metabolism in pathological heart remodeling leads to an increased reliance on glycolysis^15^, but decreased glucose/pyruvate oxidation^16^ results in a mismatch that may cause energetic defects, altered redox status, or accumulation of metabolic intermediates with signaling and physiological effects.

Many aspects of cardiac pyruvate/lactate metabolism in heart remain to be fully understood. For pyruvate to enter the mitochondrial matrix and be oxidized, it must be transported across the inner mitochondrial membrane by the mitochondrial pyruvate carrier (MPC); a hetero-oligomer composed of MPC1 and MPC2 proteins^17, 18^. Pyruvate oxidation occurs in the mitochondrial pyruvate dehydrogenase (PDH) complex and previous studies have shown that impaired cardiac PDH activity in mouse heart limits metabolic flexibility^19–22^.

However, PDH deactivation does not cause overt cardiac remodeling or dysfunction in the absence of further cardiac stress^19–22^. Another metabolic fate for pyruvate is carboxylation which is an anaplerotic reaction capable of replenishing TCA cycle intermediates. In cardiac myocytes, pyruvate carboxylation can occur in the cytosol via malic enzyme 1, or in the mitochondrial matrix via malic enzymes 2 or 3, or pyruvate carboxylase. Because MPC deletion could affect both pyruvate carboxylation and oxidation, we hypothesized that impaired MPC activity would have a greater impact on pyruvate metabolism and regulation of cardiac metabolic flexibility compared to modulating PDH activity alone.

In the present study, we demonstrate that failing human hearts express lower levels of the MPC proteins, and that loss of mitochondrial pyruvate transport and metabolism in mice is a driver of cardiac remodeling and dysfunction. Interestingly, this heart failure can be prevented or even reversed by providing mice a high-fat, low carbohydrate “ketogenic” diet. Diets with higher fat content, but enough carbohydrates to limit ketosis also significantly improved heart failure in mice lacking cardiac MPC expression. Gene expression, metabolomic analyses, and other dietary interventions all suggest improved myocardial fat metabolism, rather than increased ketone body metabolism, as the mechanism driving these improvements in heart failure. Lastly, ketogenic diet was also able to attenuate pathogenic remodeling in a surgically-induced mouse model of heart failure. These results suggest that decreased mitochondrial pyruvate metabolism induces cardiac dysfunction, and that increased dietary fat consumption may be able to prevent the fuel starvation that occurs in heart failure.

## RESULTS

### Mitochondrial Pyruvate Carrier Downregulated in Human Heart Failure

We first examined the expression of MPC proteins in heart samples of human patients obtained at the time of left-ventricular assist device (LVAD) implantation or cardiac transplantation. We compared these samples to cardiac donor tissue from hearts that were non-failing but deemed unsuitable for transplant. As expected, failing human hearts exhibited increased expression of natriuretic peptides and fibrotic collagens compared to non-failing controls (Supplemental Fig. 1a) as well as decreased expression of *PPARGC1A*, *PPARA* and mitochondrial FAO enzymes (Supplemental Fig. 1b). Failing hearts also expressed lower levels of *MPC1* and *MPC2* compared to non-failing controls at both the mRNA and protein level (Fig. 1a-c). Interestingly, failing hearts showed improvements in metabolic gene expression after LVAD placement, but natriuretic peptides and collagens were not improved (Fig. 1a-b and Supplemental Fig. 1a-b). Thus, consistent with recent data^23^, human heart failure is associated with decreased cardiac MPC expression and activity.

**Fig. 1:**
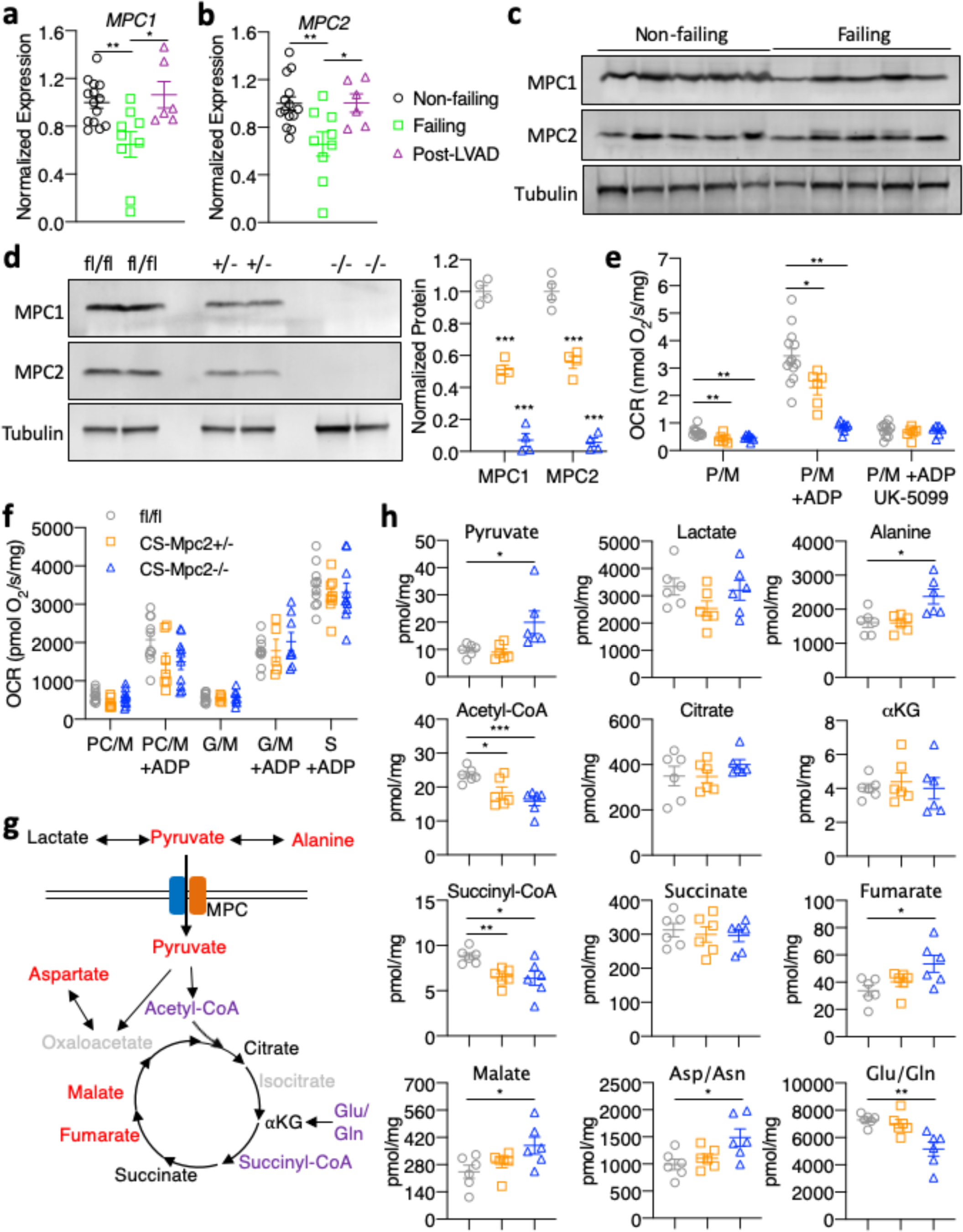
MPCs downregulated in human heart failure, deletion of cardiac MPC2 results in TCA cycle dysfunction. **a-b**, Gene expression for *MPC1* and *MPC2* normalized to *RPLP0* from human hearts of non-failing, failing, and failing hearts Post-LVAD (n=6-14). **c,** Western blot images for MPC1, MPC2, and *α*Tubulin in non-failing and failing human heart tissue (n=5). **d**, Representative western blots of MPC1, MPC2, and *α*Tubulin of mouse heart tissue and densitometry quantification (n=4). **e,** Oxygen consumption rates stimulated by pyruvate/malate (P/M) of isolated cardiac mitochondria before and after addition of ADP and 5μM of the MPC-inhibitor UK-5099 (n=6-10). **f,** Oxygen consumption rates stimulated by palmitoyl carnitine/malate (PC/M), glutamate/malate (G/M) or succinate (S) before or after the addition of ADP measured from isolated cardiac mitochondria (n=6-10). **g,** Schematic of TCA cycle alterations measured by metabolomic analyses of heart tissue. red=increased, purple=decreased, black=unchanged (comparing fl/fl to CS-Mpc2-/-), and grey=unmeasured. **h,** TCA cycle intermediates (Pyruvate, Lactate, Alanine, Acetyl-CoA, Citrate, *α*-ketoglutarate, Succinyl-CoA, Succinate, Fumarate, Malate, Aspartate/Asparagine, and Glutamate/Glutamine) measured by mass-spectrometry from 6-week old heart tissue (n=6). Mean ± s.e.m. shown within dot plot. Each symbol represents an individual sample. Two-tailed unpaired Student’s *t* test. \**P* < 0.05, \*\**P* < 0.01, \*\*\**P* < 0.001.

### CS-MPC2-/- Mice Display Altered Mitochondrial Pyruvate Metabolism and TCA Cycle Defects

To determine whether this decrease in cardiac MPC expression was an adaptive process in heart failure or contributes to the cardiac remodeling and dysfunction, we generated cardiac-specific *Mpc2* knockouts (CS-MPC2-/-) using our established *Mpc2* floxed mouse^24–26^ and mice expressing Cre under the endogenous myosin light chain 2v promoter. CS-MPC2-/- mice had complete loss of cardiac *Mpc2* gene expression (Supplemental Fig. 1c). Loss of MPC2 led to destabilization of MPC1 protein as well, and neither MPC2 nor MPC1 protein was detected in CS-MPC2-/- mouse heart (Fig. 1d). CS-MPC2-/- heart mitochondria displayed drastically reduced pyruvate stimulated oxygen consumption rates (OCR), and were resistant to inhibitory effects of the MPC inhibitor UK-5099 on respiration (Fig. 1e). *Mpc2* flox heterozygotes expressing Cre (CS-MPC2+/- mice) displayed a ∼50% decrease in MPC expression and pyruvate-stimulated respiration (Fig. 1d-e and Supplemental Fig. 1c). Isolated mitochondria from CS-MPC2+/- and CS-MPC2-/- hearts displayed normal OCR on palmitoylcarnitine/malate, glutamate/malate, and succinate (Fig. 1f), suggesting a specific defect in mitochondrial pyruvate metabolism. CS-MPC2-/- mice also displayed slight, but significantly elevated blood lactate levels (Supplemental Fig. 1d), consistent with the heart as an appreciable lactate-consuming organ.

To more thoroughly investigate how loss of MPC expression altered mitochondrial metabolism, targeted metabolomics analyses for organic acids, amino acids, short chain acyl-CoAs, and acylcarnitines were conducted with hearts from 6-week old female mice (Fig. 1g-h and Supplemental Table 1). As summarized in Fig. 1g, CS-MPC2-/- hearts contained decreased acetyl-CoA levels, and an accumulation of TCA cycle intermediates upstream of acetyl-CoA (fumarate, malate, and oxaloacetate [aspartate measured as surrogate])(Fig. 1g-h). Altogether, these findings suggest that loss of cardiac MPC results in defective mitochondrial pyruvate metabolism and alterations in TCA cycle flux.

### CS-MPC2-/- Mice Develop Dilated Cardiomyopathy

Hearts from 6-week old CS-MPC2-/- mice appeared normal by echocardiography, but heart weight and hypertrophic gene expression was slightly elevated in these young mice (Fig. 2a-c, Supplemental Fig. 1e-g, and Supplemental Fig. 2a-h). Cardiac enlargement and decreased contractile function was well-evident at 10-weeks and further worsened at 16-weeks of age (Fig. 2a-d and Supplemental Fig. 2a-h). Increased ventricular mass was confirmed at sacrifice (Fig. 2e-f), as was increased lung weight indicative of lung edema (Fig. 2g). CS-MPC2-/- hearts also showed dramatically altered gene expression markers of heart failure and fibrosis (Fig. 2h-i).

**Fig. 2:**
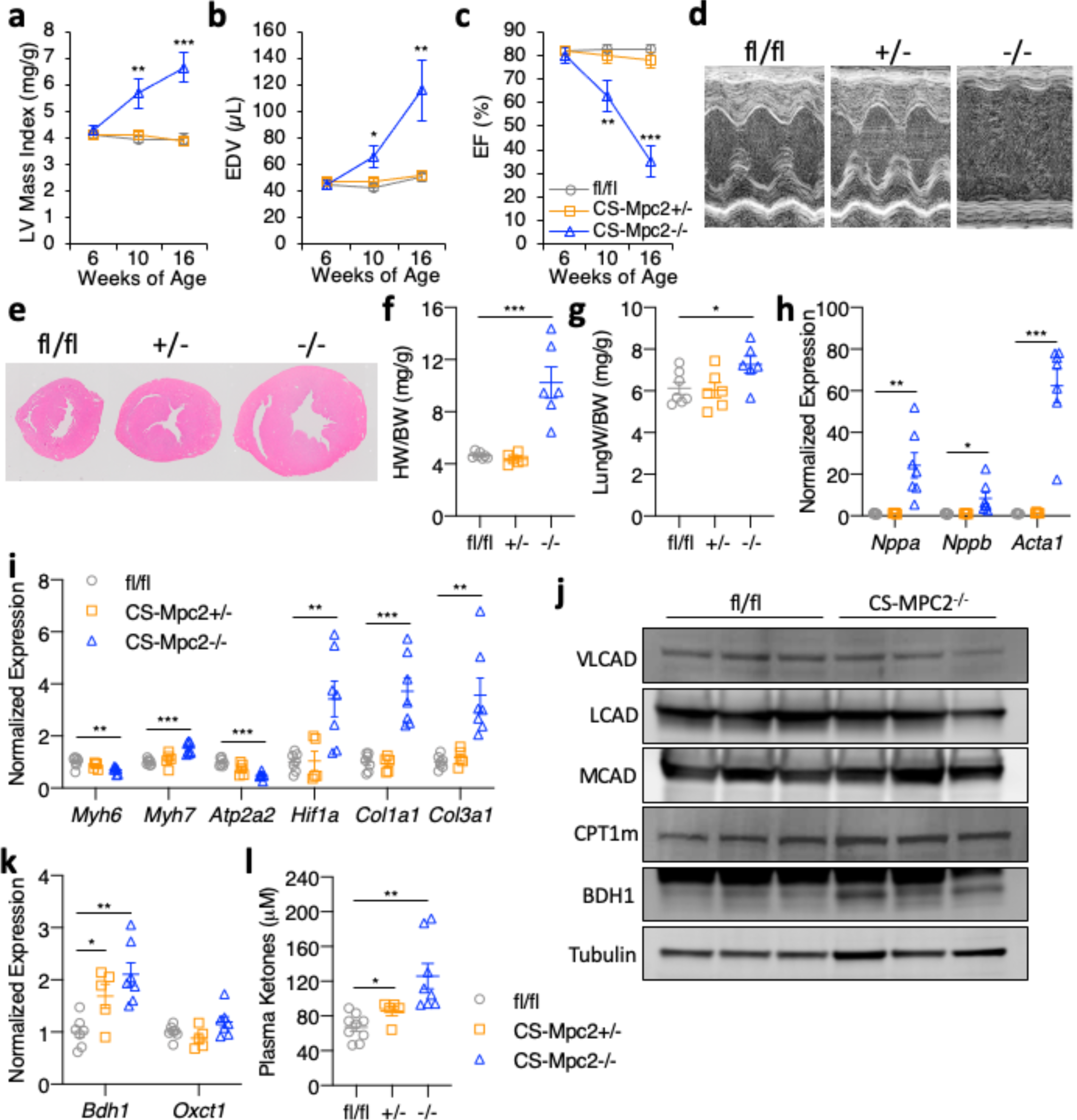
CS-MPC2-/- mice develop dilated cardiomyopathy. **a-c,** Echocardiography measures of left ventricular (LV) mass index, end-diastolic volume (EDV), and ejection fraction (EF) of mice at 6, 10, and 16-weeks of age (n=7-10). **d,** Representative M-mode electrocardiogram images of 16-week old mice. **e,** Representative short-axis heart images stained by H&E. **f-g,** Heart weight and lung weight normalized to body weight (n=6). **h-i,** Gene expression markers of cardiac hypertrophy/failure from 16-week old mouse hearts (n=6). **j,** Western blot images of VLCAD, LCAD, MCAD, CPT1B, BDH1, and *α*Tubulin from whole cardiac lysate (n=3). **k,** Gene expression for *Bdh1* and *Oxct1* from 16-week old mouse hearts (n=6). **l,** Plasma total ketone body levels from 16-week old mice (n=6). Mean ± s.e.m. shown within dot plot. Each symbol represents an individual sample. Two-tailed unpaired Student’s *t* test. \**P* < 0.05, \*\**P* < 0.01, \*\*\**P* < 0.001.

Importantly, CS-MPC2+/- heterozygotes displayed normal cardiac size, function, and hypertrophic gene expression (Fig. 2a-i and Supplemental Fig. 2a-h), suggesting that MPC haploinsufficiency or Cre expression alone was not provoking this heart failure phenotype. WT C57BL6/J mice treated with the MPC inhibitor MSDC-0602, currently in development to treat diabetes and nonalcoholic steatohepatitis^26^, also did not show cardiac enlargement or cardiac hypertrophic gene expression (Supplemental Fig. 2i-j). Together, these results indicate that complete loss of MPC expression, but not partial loss or pharmacologic MPC inhibition, results in cardiac remodeling and dysfunction.

Interestingly, other than *Cpt1b*, these CS-MPC2-/- hearts did not show the downregulation of PPAR*α* target gene expression such as enzymes and transporters associated with FAO as is typical for failing hearts (Fig. 2j and Supplemental Fig. 2k). CS-MPC2-/- hearts exhibited increased expression of BDH1 at the gene and protein level (Fig. 2j-k) and significantly elevated plasma ketone bodies were found in CS-MPC2-/- mice (Fig. 2l). Together, this elevated ketosis and increased BDH1 expression suggests increased ketone body metabolism in the failing CS-MPC2-/- hearts^27, 28^, which was recently shown to be an adaptive and protective process in heart failure^29^.

### Ketogenic Diet Fully Prevents Heart Failure in CS-MPC2-/- Mice

Since CS-MPC2-/- hearts appeared to have maintained capacity for FAO as well as potentially increased ketone body utilization, we hypothesized that the cardiac remodeling and dysfunction in CS-MPC2-/- mice could be improved by providing nutrients in the diet that could be used, and removing those that could not. To test this, fl/fl and CS-MPC2-/- mice were fed a low-carbohydrate (1.8% kcal), high-fat (93.9% kcal) “ketogenic diet” (KD) or a low-fat (LF) control diet from 6 weeks until 17 weeks of age. KD resulted in the expected increase in ketosis (Fig. 3a), as well as limited weight gain, decreased blood glucose, and decreased plasma insulin concentrations in both fl/fl and CS-MPC2-/- mice (Supplemental Fig. 3a-c). LF-fed CS-MPC2-/- mice displayed extreme cardiac enlargement and dysfunction (Fig. 3b-d and Supplemental Fig. 3d-n), which was even worse when compared to chow-fed CS-MPC2-/- mice (see Fig. 2), potentially due to the increased content of refined sucrose in the LF diet. Strikingly, KD-fed CS- MPC2-/- mice displayed virtually normal cardiac size and function during echocardiography studies at 10- and 16-weeks of age (Fig. 3b-d, Supplemental Fig. 3d-n, and Supplemental Video 1). The severe cardiac dysfunction in LF-fed CS-MPC2-/- mice was associated with loss of body weight by 17 weeks of age (Supplemental Fig. 3a), which was driven by loss of adipose tissue fat mass (Supplemental Fig. 3o-s). Nearly 35% of LF-fed CS-MPC2-/- mice died prior to 17 weeks of age, but all CS-MPC2-/- mice fed KD survived (Fig. 3e). Extreme LV dilation, cardiac enlargement, and increased lung edema was evident in LF-fed CS-MPC2-/- mice at sacrifice, which was completely prevented by feeding KD (Fig. 3f-h). Gene expression markers for heart failure and fibrosis, as well as trichrome fibrosis staining, were all significantly altered in LF-fed, and completely corrected in KD-fed, CS-MPC2-/- hearts (Fig. 3h-n). LF-fed CS-MPC2-/- hearts also displayed altered hypertrophic growth signaling by increased ERK phosphorylation, decreased AMPK*α* phosphorylation, increased mTOR phosphorylation, and increased S6 ribosomal protein phosphorylation (Fig. 3o), consistent with increased protein synthesis required to drive pathologic cardiac hypertrophy^30^. Feeding CS-MPC2-/- mice KD completely prevented this aberrant hypertrophic growth signaling (Fig. 3o). Altogether, these results show that ketogenic diet is able to completely prevent cardiac remodeling and dysfunction of CS-MPC2-/- mice.

**Fig. 3:**
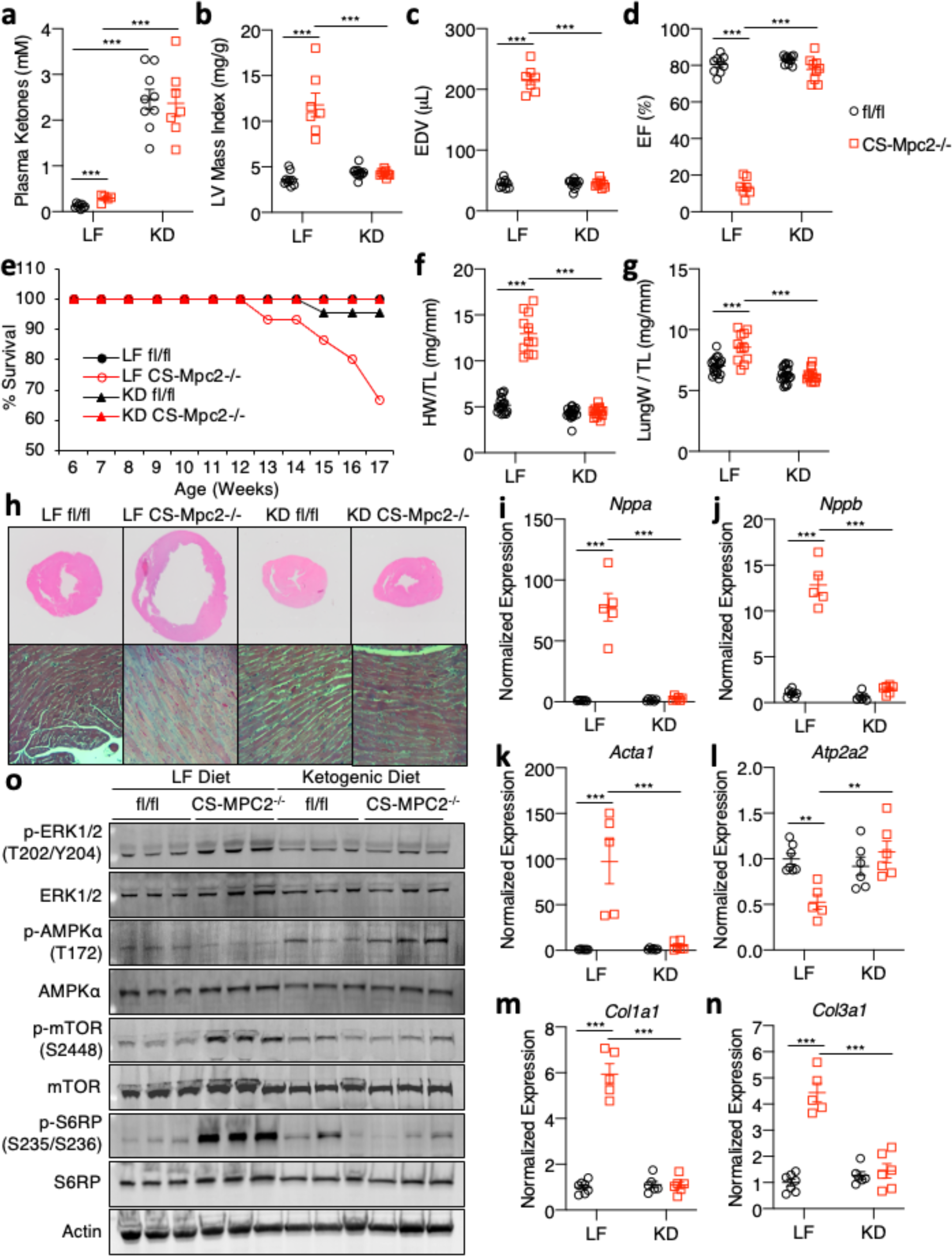
Ketogenic diet can prevent heart failure in CS-MPC2-/- mice. **a,** Plasma total ketone bodies from low fat (LF)- or ketogenic diet (KD)-fed mice (n=5-9). **b-d,** Echocardiography measures of left ventricular (LV) mass index, end-diastolic volume (EDV), and ejection fraction (EF) of LF- or KD-fed mice at 16-weeks of age (n=7-11). **e,** Survival curve of LF- or KD-fed mice (initial n=14-20). **f**-**g,** Heart weight and lung weight normalized to tibia length of LF- or KD-fed 17-week old mice (n=11-20). **h,** Representative short-axis H&E images and magnified trichrome stains of hearts from LF- or KD-fed mice. **i-n,** Gene expression markers of cardiac hypertrophy/failure and fibrosis from mouse hearts (n=5-7). **o,** Western blot images for signaling pathways associated with cardiac hypertrophic growth (PhosphoERK, Total ERK, PhosphoAMPK*α*, Total AMPK*α*, Phospho-mTOR, Total mTOR, Phospho-S6-Ribosomal Protein, Total S6-Ribosomal Protein, and *β*-Actin) from hearts of LF- or KD-fed mice (n=3). Mean ± s.e.m. shown within dot plot. Each symbol represents an individual sample. Two-way ANOVA with Tukey’s multiple-comparisons test. \**P* < 0.05, \*\**P* < 0.01, \*\*\**P* < 0.001.

### Ketogenic Diet Downregulates Cardiac Ketone Body Oxidation and Enhances Fatty Acid Metabolism

The chow-fed CS-MPC2-/- mice showed elevated ketone bodies and increased BDH1 expression (Fig. 2j-l), consistent with recent work suggesting an increase in ketone body oxidation in failing hearts^27, 28^. LF-fed CS-MPC2-/- mice also displayed an increase in plasma ketone bodies (Fig. 3a), and the failing hearts from these mice showed an upregulation of the ketolytic enzymes *Bdh1* and *Oxct1*, as well as increases in C4-OH-carnitine and 3- hydroxybutyrate-CoA (Fig. 4a-f). Interestingly, hearts from both fl/fl and CS-MPC2-/- mice show decreased BDH1 and *Oxct1* expression after KD-feeding (Fig. 4b-d), suggesting hearts downregulate ketone body oxidation during ketogenic diet^31^. Along these lines, the levels of succinyl-CoA, succinate, and succinate/succinyl-CoA ratio all suggest increased ketolytic flux in failing LF-fed CS-MPC2-/- hearts that is counter-intuitively reduced by KD-feeding (Fig. 4g-i). KD-feeding also normalized the levels of free CoA-SH, malonyl-CoA, as well as the expression of malonyl-CoA-generating enzymes *Acaca* and *Acacb* in CS-MPC2 hearts (Fig. 4j-n). Altogether, these results suggest that the improvements in cardiac remodeling and function from KD-feeding were not related to ketone metabolism.

**Fig. 4:**
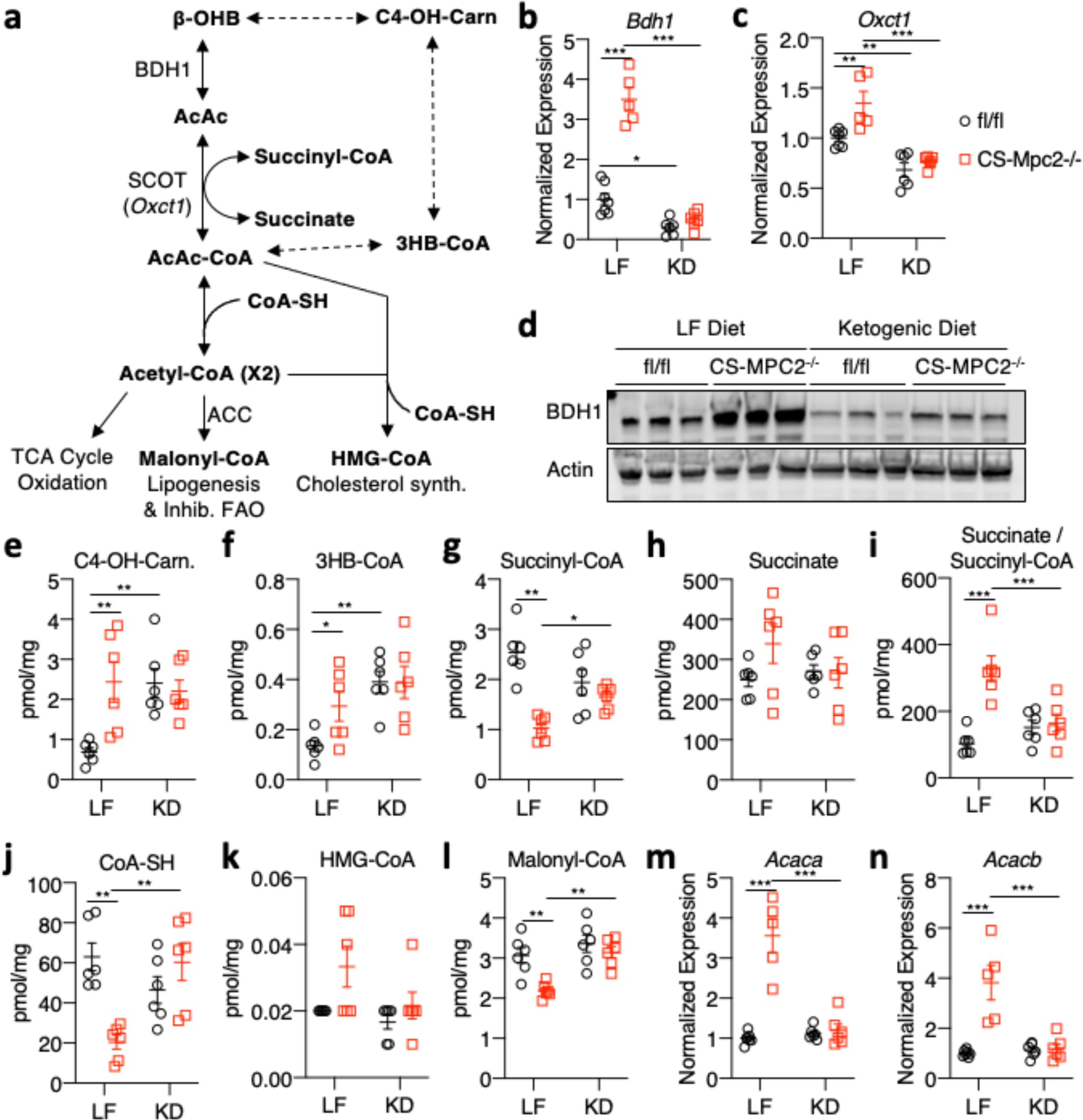
Ketogenic diet downregulates cardiac ketone body catabolism. **a,** Schematic of oxidative and non-oxidative ketone body catabolism. **b**-**c,** Gene expression for the ketolytic enzymes *Bdh1* and *Oxct1* from hearts of low fat (LF)- or ketogenic diet (KD)-fed fl/fl or CS- Mpc2-/- mice (n=5-7). **d,** Western blot images of BDH1 and Actin from heart tissue of LF- or KD-fed mice (n=3). **e-l,** Cardiac concentrations of metabolites associated with ketone body catabolism measured in hearts from LF- or KD-fed mice (n=6). **m-n,** Gene expression for *Acaca* and *Acacb* normalized to *Rplp0* from hearts of LF- and KD-fed mice (n=5-7). Mean ± s.e.m. shown within dot plot. Each symbol represents an individual sample. Two-way ANOVA with Tukey’s multiple-comparisons test. \**P* < 0.05, \*\**P* < 0.01, \*\*\**P* < 0.001.

The failing LF-fed CS-MPC2-/- hearts displayed an accumulation of acylcarnitines and depletion of free carnitine, which was normalized by KD-feeding (Fig. 5a-d, and Supplemental Table 2). Accumulation of acylcarnitines suggests a decrease in their transport into the mitochondrial matrix and oxidation. Indeed, failing LF-fed CS-MPC2-/- hearts displayed decreased expression of *Ppargc1a*, *Ppara*, and many of their target genes for fatty acid transport and metabolism (Fig. 5e-l). KD-feeding rescued or strongly elevated the expression of *Ppargc1a*, *Ppara*, and its gene targets related to FAO in both fl/fl and CS-MPC2-/- hearts (Fig. 5e-l).

**Fig. 5:**
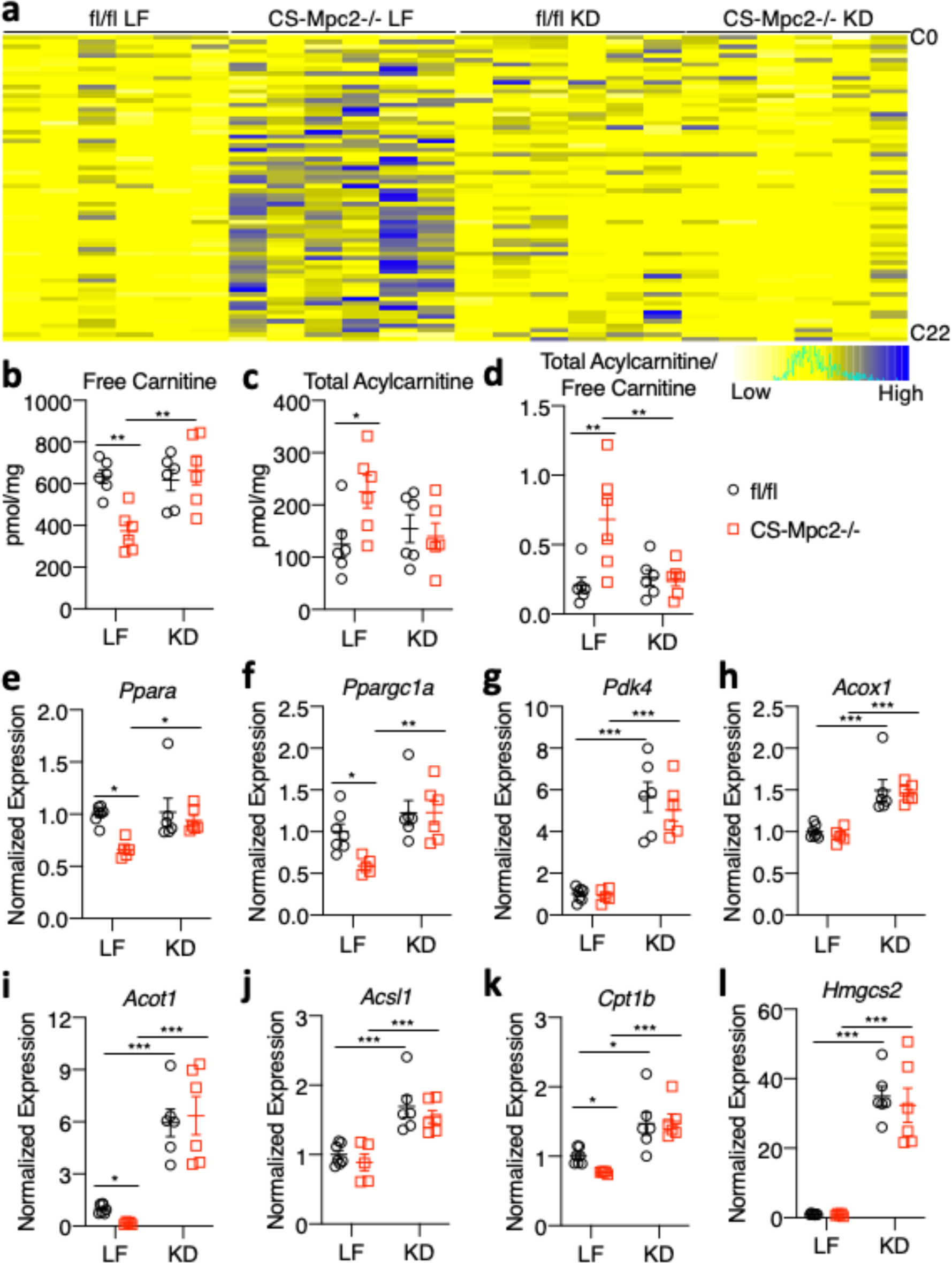
Ketogenic diet enhances cardiac fatty acid metabolism. **a,** Heatmap of acylcarnitine species measured in hearts of low fat (LF)- or ketogenic diet (KD)-fed fl/fl or CS-Mpc2-/- mice (n=6). **b-d,** Concentrations of free carnitine, total acylcarnitines, and the acylcarnitine/free carnitine ratio measured by mass-spectrometry of heart tissue (n=6). **e-l,** Gene expression markers of PPAR*α* and fatty acid oxidation (*Ppara*, *Ppargc1a*, *Pdk4*, *Acox1*, *Acot1*, *Acsl1*, *Cpt1b*, and *Hmgcs2*) from heart tissue of LF- or KD-fed mice (n=5-7). Mean ± s.e.m. shown within dot plot. Each symbol represents an individual sample. Two-way ANOVA with Tukey’s multiple-comparisons test. \**P* < 0.05, \*\**P* < 0.01, \*\*\**P* < 0.001.

Interestingly, the *Ppara* target gene, *Hmgcs2,* which generates ketone bodies and is normally expressed almost exclusively in the liver, was strongly induced in KD-fed fl/fl and CS-MPC2-/- hearts (Fig. 5l). Cumulatively, these results suggest that KD-feeding does not enhance cardiac ketone body metabolism, but rather stimulates FAO, which may be responsible for the improved cardiac remodeling and performance.

### Exogenous Ketone Bodies Moderately Attenuate Cardiac Remodeling in CS-MPC2-/- Mice

We also wanted to assess whether increased ketosis without altering dietary fat intake was able to improve cardiac function. To test this, CS-MPC2-/- mice were maintained on chow diet and injected i.p. with saline vehicle or 10 mmol/kg *β*-hydroxybutyrate (*β*HB) daily for two weeks (Supplemental Fig. 4a). Over this timeframe, vehicle treated CS-MPC2-/- mice displayed worsened LV dilation and contractile function, which were either limited or improved by daily *β*HB administration (Supplemental Fig. 4b-h). At sacrifice, 4 hours after the last *β*HB injection, plasma ketone concentrations were significantly elevated (Supplemental Fig. 4i), but not nearly to the same degree as when fed a ketogenic diet (see Fig. 3a). Heart weight (Supplemental Fig. 4j) and hypertrophic/fibrotic gene expression (Supplemental Fig. 4k) were only modestly improved by administering ketone bodies daily on top of carbohydrate-rich chow diet.

In a second attempt to raise ketosis without altering dietary fat, we fed mice a diet supplemented with 16.5%kcal ketone esters (KE). For this experiment, mice were fed control or KE diet from 9-15 weeks of age. KE diet slightly raised plasma ketone bodies (Supplemental Fig. 5a), but did not improve cardiac size or contractile function measured by echocardiography (Supplemental Fig. 5b-e), or heart weight at sacrifice (Supplemental Fig. 5f). Lastly, cardiac gene expression of markers of heart failure were only modestly improved by KE diet (Supplemental Fig. 5g-i). Thus, two different ways to enhance ketosis without altering dietary fat intake did not drastically improve the cardiac size and function of CS-MPC2-/- mice. These results suggest that other factors are the predominant driver for improving heart failure in KD-fed CS-MPC2-/- mice.

### High-Fat Diets Significantly Improve Heart Failure in CS-MPC2-/- Mice

To dissect the importance of dietary fat and myocardial FAO, we also fed fl/fl and CS- MPC2-/- mice two diets that were higher in fat, but with moderate levels of carbohydrate and protein, which failed to, or only modestly increased, plasma ketone body concentrations (Fig. 6a-b). Feeding CS-MPC2-/- mice a ∼42% medium chain triglyceride (MCT) or a 60% high-fat (HF) diet was also able to significantly improve cardiac enlargement and contractile function as measured by echocardiography (Fig. 6c-d, Supplemental Fig. 6a-i, and Supplemental Video 2).

**Fig. 6:**
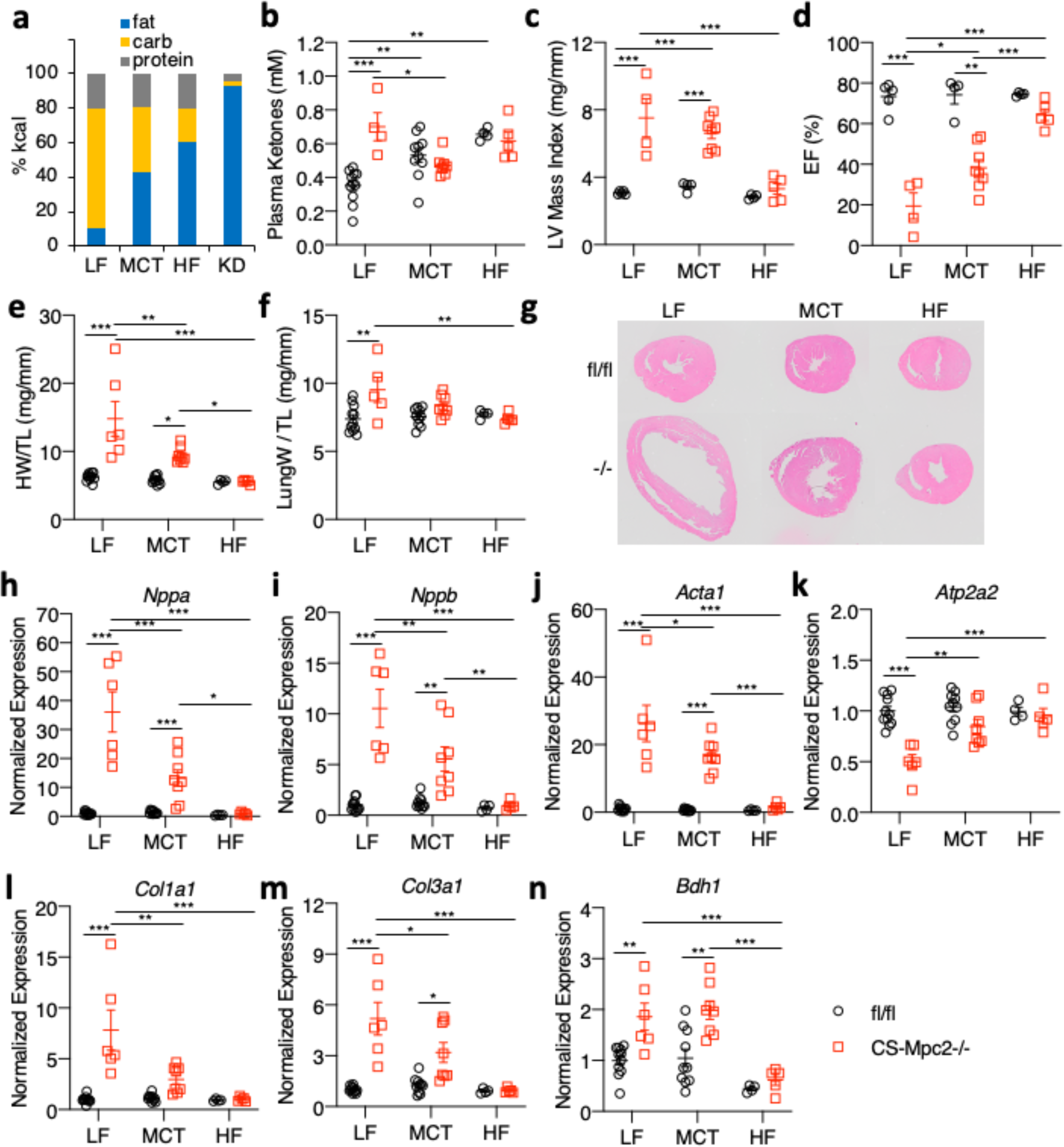
High fat diets also prevent cardiac remodeling and dysfunction in CS-MPC2-/- mice. **a,** Graphical comparison of diet macronutrient composition for low fat (LF), medium chain triglyceride (MCT), high fat (HF), and ketogenic diet (KD). **b,** Plasma total ketone body concentrations measured from mice after LF, MCT, or HF diet feeding (n=4-11). **c**-**d,** Echocardiography measures of left ventricular (LV) mass index and ejection fraction (EF) of mice fed LF, MCT, or HF diets (n=4-11). **e**-**f,** Heart weight and lung weight normalized to tibia length (n=4-11). **g,** Representative short-axis heart images stained with H&E. **h-n,** Gene expression markers of hypertrophy, heart failure, fibrosis, and the ketolytic enzyme *Bdh1* from mouse hearts (n=4-11). Mean ± s.e.m. shown within dot plot. Each symbol represents an individual sample. Two-way ANOVA with Tukey’s multiple-comparisons test. \**P* < 0.05, \*\**P* < 0.01, \*\*\**P* < 0.001.

Heart weight, lung edema, and hypertrophic/fibrotic gene expression were also significantly improved by MCT and HF diets (Fig. 6e-n). Interestingly, the HF diet was also capable of enhancing expression of *Ppara* and its target genes (Supplemental Fig. 6j-l), as well as lowering *Bdh1* expression (Fig. 6n) compared to LF diet. Thus, diets enriched with higher levels of fat but enough carbohydrate and protein to limit ketosis were also able to significantly improve or even prevent cardiac remodeling and dysfunction in CS-MPC2-/- mice.

### Ketogenic Diet Can Reverse Heart Failure in CS-MPC2-/- Mice

With the dramatic effects of KD fully preventing heart failure, we were also curious to assess if KD could reverse existing heart failure. We allowed CS-MPC2-/- mice to consume chow diet until 16 weeks of age, then assigned them to either LF- or KD-feeding for 3 weeks (Fig. 7a). All CS-MPC2-/- mice displayed cardiac dilation and poor contractile function during the 16-week echocardiograms, which was then worsened further by 3 weeks of LF diet feeding (Fig. 7b-d and Supplemental Fig. 7a-h). However, 3-weeks of KD-feeding greatly improved the LV dilation and contractile function of the previously failing CS-MPC2-/- hearts (Fig. 7b-d, Supplemental Fig. 7a-h, and Supplemental Video 3). The 3 weeks of KD-feeding strongly elevated ketosis (Fig. 7e) and heart weight, lung edema, and cardiac gene expression markers of pathological remodeling and fibrosis were all drastically reversed by 3-weeks of KD-feeding (Fig. 7f-h). Thus, ketogenic diet consumption for only 3-weeks was capable of producing “reverse remodeling” of the heart failure observed in CS-MPC2-/- mice.

**Fig. 7:**
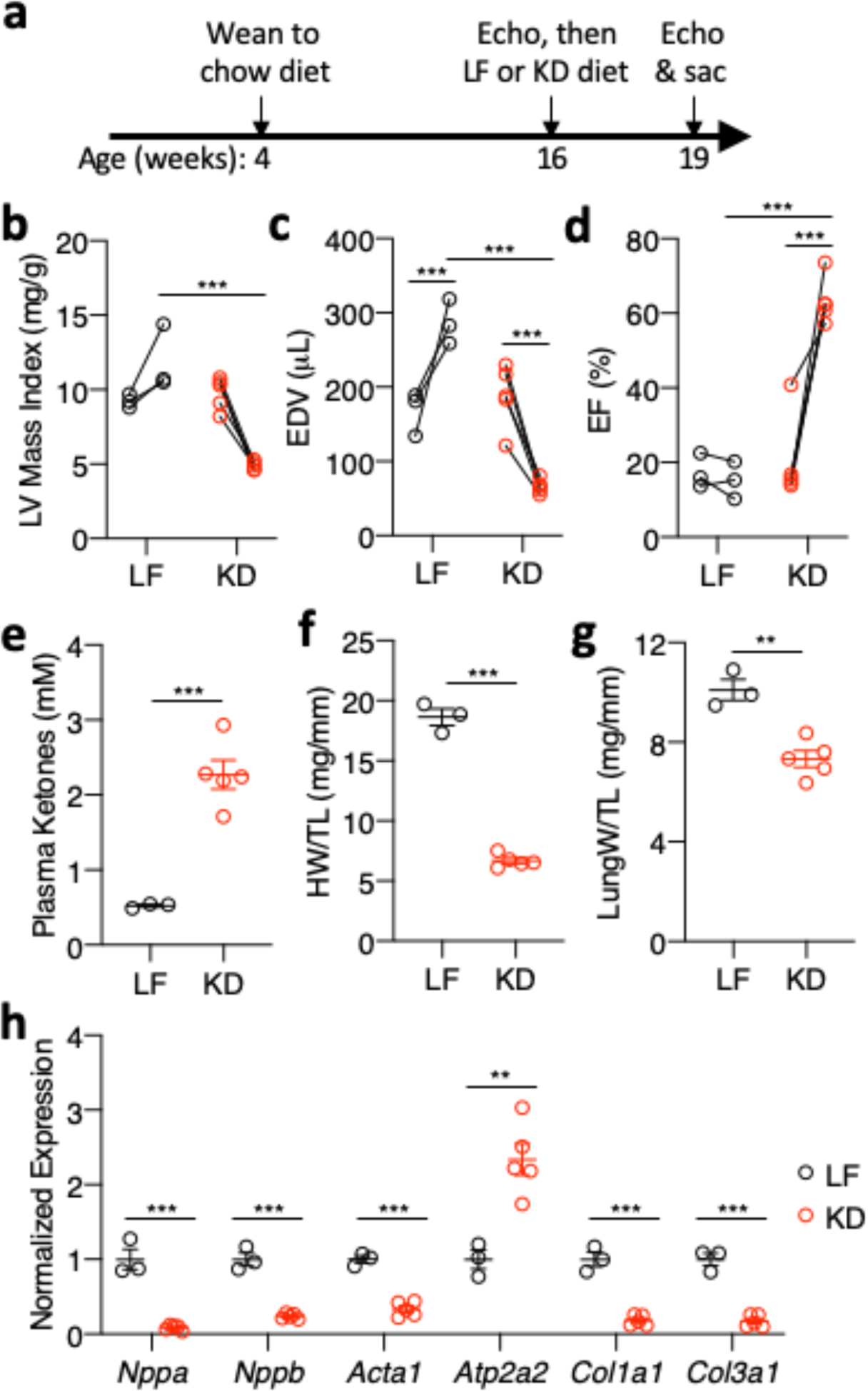
Ketogenic diet can reverse heart failure in CS-Mpc2-/- mice. **a,** Timeline for heart failure reversal experiment, in which CS-MPC2-/- mice were switched to low fat (LF) or ketogenic diet (KD) at 16 weeks of age for 3 weeks. **b-d,** Echocardiography measures of left ventricular (LV) mass index, end-diastolic volume (EDV), and ejection fraction (EF) of CS- MPC2-/- mice PRE and POST LF or KD feeding (n=3-5; data presented as PRE-POST with first data point at 16-weeks old and second data point at 19-weeks old after 3 weeks of LF or KD). **e,** Plasma total ketone values from CS-Mpc2-/- mice fed LF or KD (n=3-5). **f**-**g,** Heart weight and lung weight normalized to tibia length (n=3-5). **h,** Gene expression markers of hypertrophy, heart failure, and fibrosis from mouse hearts (n=3-5). Data presented either as PRE-POST, or mean ± s.e.m. shown within dot plot. Each symbol represents an individual sample. Two-tailed unpaired Student’s *t* test. \**P* < 0.05, \*\**P* < 0.01, \*\*\**P* < 0.001.

### Ketogenic Diet Decreases Cardiac Remodeling in an Ischemic, Pressure-Overload Model of Heart Failure

As KD was able to prevent and reverse heart failure in CS-MPC2-/- mice and MPC expression is reduced in human heart failure, we wondered if KD would also improve heart failure in induced models of pathological remodeling in mice. We subjected WT C57BL/6J mice to combined transverse aortic constriction plus apical myocardial infarction (TAC+MI) surgery which results in consistent LV dilation and failure^32^, and led to a downregulation of the *Mpc1* and *Mpc2* genes (Fig. 8a). Two weeks after TAC+MI surgery, mice were imaged by echocardiography and randomized to LF or KD for the following 2 weeks (Fig. 8b). Two weeks of KD was sufficient to induce robust ketosis, and drastically decrease blood glucose and plasma insulin values (Supplemental Fig. 8a-c). All mice displayed cardiac dilation and decreased contractility 2 weeks post TAC+MI (Fig. 8c-e and Supplemental Fig. 8d-e). While LF-fed mice displayed further cardiac remodeling and enlargement 4 weeks post TAC+MI, this was prevented by KD-feeding despite an identical aortic pressure gradient (Fig. 8c-e, Supplemental Fig. 8d-e, and Supplemental Video 4). Contractile function measured by ejection fraction (EF) was greatly reduced two weeks post TAC+MI and trended towards being worsened with 2 weeks of LF-feeding but not KD-feeding (Fig. 8e). Lastly, KD significantly reduced or provided strong trends towards reduction in heart weight, lung edema, and hypertrophic gene expression (Fig. 8f-i) in this TAC+MI model of dilated cardiomyopathy. These data suggest that KD attenuates cardiac remodeling and dysfunction from the TAC+MI model of heart failure, similar to as previously reported^29^.

**Fig. 8:**
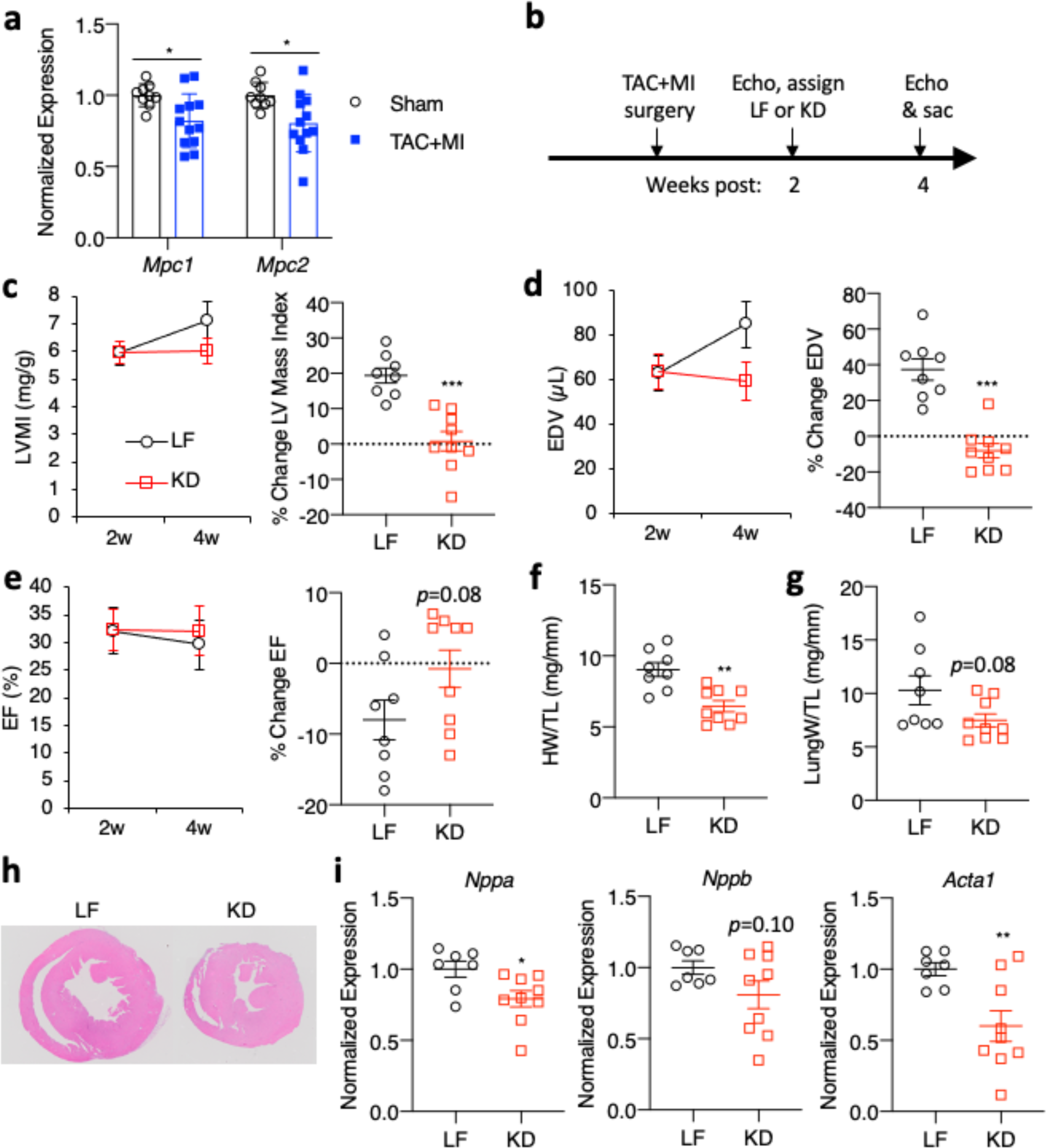
Ketogenic diet reduces cardiac remodeling from TAC+MI ischemic and pressure-overload induced heart failure. **a,** Gene expression for *Mpc1* and *Mpc2* from hearts of wildtype C57BL6/J mice after sham or transverse aortic constriction plus myocardial infarction (TAC+MI) surgery (n=9-12). **b,** Timeline for TAC+MI study in which wildtype C57BL6/J mice are switched to low fat (LF) or ketogenic diet (KD) 2 weeks post-surgery and fed the diets for 2 weeks prior to sacrifice. **c-e,** Echocardiography measures of LV mass index, EDV, and EF, along with the %-change between echocardiography at 2- and 4-weeks post-surgery (n=8-9). **f**-**g,** Heart weight and lung weight normalized to tibia length of mice subjected to TAC+MI surgery (n=8- 9). **h,** Representative short axis H&E heart images after TAC=MI surgery. **i,** Gene expression of *Nppa*, *Nppb*, and *Acta1* from hearts subjected to TAC+MI surgery and LF or KD feeding (n=6). Data are presented as mean ± s.e.m. within dot plot. Each symbol in dot plot represents an individual sample. Two-tailed unpaired Student’s *t* test. \**P* < 0.05, \*\**P* < 0.01, \*\*\**P* < 0.001.

## DISCUSSION

An estimated 6.2 million American adults have heart failure and this number is increasing due to the aging population and prevalence of cardiovascular risk factors^33^. Myocardial fuel metabolism is altered in hypertrophy and heart failure, characterized as a generalized decrease in the ability to oxidize fatty acids and other substrates in the mitochondrion^15, 34^. While glycolysis is increased^35–37^, this may not be sufficient to compensate for diminished mitochondrial metabolism of pyruvate and fats resulting in the metabolically “starved” failing heart. Combating the metabolic remodeling that occurs in heart failure is a tempting target for therapeutic intervention.

The import of pyruvate into the mitochondria occurs via the mitochondrial pyruvate carrier, which was identified in 2012 as a hetero-oligomeric complex of MPC1 and MPC2 proteins^17, 18^. An early study conducted prior to the cloning of MPC proteins and using a chemical inhibitor estimated that cardiac MPC expression was quite high, and MPC activity would be rate-limiting for pyruvate oxidation in heart mitochondria^38^. However, studies regarding the importance of MPC activity in cardiac function or development of heart failure have been quite limited. Expression of MPC1 and MPC2 was shown to be an important marker of surviving myocardium near the border of infarct zones in a pig model, and this study also identified increased MPC expression in human hearts with ischemic heart failure^39^. While this current work was in preparation, another report showed that failing human hearts exhibited decreased expression of the MPC proteins^23^, which we have confirmed in this current study. Thus, myocardial MPC expression in heart failure may depend on ischemic vs non-ischemic etiology, as well as location in relation to infarct zone. Together with two companion papers, we also now show that complete deletion of MPC in myocardium leads to a severe, progressive cardiac remodeling and dilated heart failure. However, pharmacologic MPC inhibition or loss of one MPC2 allele and approximately 50% of the MPC protein did not affect cardiac function. This likely highlights the differences between the heart failure models used in these studies. While heart failure caused by hemodynamic stressors in mice and humans reduced *Mpc* expression by ∼30% (Fig. 1 and Fig. 8), heterozygous CS-MPC2+/- mice were completely normal in the absence of cardiac stress (Fig. 2). These findings suggest that partial inhibition of MPC activity in the heart is not limiting and can be overcome metabolically, as long as other cardiac stressors are not restricting metabolic flexibility.

Previous work has shown that modulating the expression or activity of PDH limits cardiac metabolic flexibility by decreasing glucose oxidation and increasing FAO^19–22^.

Interestingly, these models of decreased PDH activity did not result in overt cardiac dysfunction. One possible explanation for why MPC-deletion is more severe is that blocking pyruvate entry could also impact pyruvate carboxylation (anaplerosis) and the replenishing of TCA cycle intermediates. Although the effects of deleting pyruvate carboxylase in the myocardium are not known, this pathway is known to be active in the heart^40^. However, the majority of pyruvate carboxylation in the heart likely occurs by NADP+-dependent malic enzyme^41^ generating malate in the cytosol. Therefore, the severe cardiac dysfunction and remodeling in CS-MPC2-/- hearts may not be due to loss of combined pyruvate oxidation and carboxylation.

The current studies cannot definitively explain why CS-MPC2-/- mice develop heart failure. The simplest explanation of an inability to oxidize pyruvate resulting in an energetic deficit is possible. But with normal adult hearts deriving a high percentage of their ATP from FAO^34^, perhaps additional mechanisms contribute to the heart failure in CS-MPC2-/- mice.

Another possibility is that a decrease in mitochondrial pyruvate metabolism results in a metabolic mismatch and an accumulation of a metabolic intermediate that enhances hypertrophic signaling. An example of this would be the oncometabolite 2-hydroxyglutarate (2-HG), which has been implicated in driving cardiac hypertrophy and impairing contractility^42, 43^. Interestingly, we found that failing LF-fed CS-MPC2-/- hearts contained almost 2-fold higher concentrations of total 2-HG (Supplemental Table 2). However, hearts from KD-fed mice also had higher total 2-HG compared to LF-fed fl/fl mice (Supplemental Table 2). Unfortunately, our mass-spectrometry analyses did not distinguish between D- and L-2-HG, as only D-2-HG appears to be responsible for inducing cardiomyopathy^42, 43^. Another theory is that a decrease in glucose oxidation could result in a shift to accessory glucose utilization pathways. Importantly, these accessory pathways can trigger lipid or post-translational protein modifications such as glycosylation, O-GlcNAcylation, or advanced glycation end products, which have all been implicated in heart failure^44–48^. Altered signaling or protein/lipid modifications as a result of decreased glucose oxidation in heart failure are exciting hypotheses that certainly warrant further investigation.

One of the most striking findings of this work is that feeding a low carbohydrate, high fat “ketogenic diet” completely prevented or reversed heart failure in CS-MPC2-/- mice. Several lines of evidence suggest that these improvements were driven primarily by enhanced provision of fatty acids rather than ketone bodies. While hearts can extract and metabolize ketone bodies in proportion to their delivery, ketones and fatty acids are in competition for oxidation^5–7^. In agreement with a previous report^31^, we show that hearts from mice fed a ketogenic diet decrease the expression of the ketolytic enzymes BDH1 and *Oxct1*, and rely more on FAO. This KD-feeding was also associated with upregulation of PPAR*α*-target genes related to FAO and corrected the cardiac accumulation of acylcarnitines. Lastly, diets that were enriched with fat, but not overly ketogenic due to moderate levels of carbohydrate and protein, were also able to significantly prevent heart failure in CS-MPC2-/- mice. It is interesting that the degree of heart failure improvement appears to track with the amount of dietary fat and likewise reduction of dietary carbohydrate. Along these lines, it is also interesting that hearts from CS-MPC2-/- mice showed even worse failure after refined LF diet feeding compared to chow (Fig. 3 and Supplemental Fig. 3 compared to Fig. 2 and Supplemental Fig. 2), potentially due to the large amount of sucrose in the LF diet compared to complex carbohydrates in chow. Therefore, we believe enhanced FAO and limiting the provision of carbohydrate to be the predominant mechanism for improved cardiac function in CS-MPC2-/- mice.

We also attempted to also raise ketosis without modulating dietary fat. Injecting CS- MPC2-/- mice daily with *β*-hydroxybutyrate did slightly ameliorate cardiac remodeling, but feeding a ketone-ester-supplemented chow did not improve cardiac size or function. Recent studies have described improvements in cardiac function with ketone body infusion in both a dog model and human patients with heart failure^29, 49^. Additionally, mouse models of BDH1 loss or overexpression or OXCT1 loss suggest that increased ketone metabolism is a protective adaptation in heart failure^29, 50, 51^. A limitation of these models we used is that the circulating ketone concentrations generated by either method are not as high as when feeding a ketogenic diet. Thus, it is difficult to say whether a more pronounced level of ketosis would also improve these CS-MPC2-/- hearts.

Lastly, KD was also able to attenuate cardiac remodeling in a TAC+MI pressure overload mouse model of heart failure, in agreement with a recent report^29^. KD has previously been used to improve heart failure in a mouse model of primary mitochondrial oxidative phosphorylation defects as well^52^. And several case reports have described ketogenic diets to improve cardiomyopathy in subjects with glycogen storage disease type III^53–56^. Interestingly, a ketogenic diet was unable to improve cardiac hypertrophy in a mouse model of defective FAO caused by carnitine palmitoyltransferase 2 deletion^57^. This further suggests that ketogenic diet may improve heart failure predominantly by increasing myocardial FAO rather than elevating cardiac ketone oxidation. Additionally, there is substantial literature describing improvement in rodent cardiomyopathy models using “non-ketogenic” high fat or PUFA-enriched diets in the absence of obesity^58^. Interestingly, ketogenic diet was recently shown to improve cardiac remodeling in the obese *db/db* mouse model^59^, potentially due to the improved insulin sensitivity with significant carbohydrate restriction. However, in the setting of obesity and insulin resistance, increased dietary fat may add to cardiac lipid accumulation or “lipotoxicity”^60^. How dietary fat improves cardiomyopathy, and whether it counteracts the loss of PPAR*α* in heart failure are future questions. It is also interesting to speculate that the reduction in blood glucose and resulting increase in plasma ketones is responsible for the reduced heart failure mortality seen with sodium-glucose transporter 2 (SGLT2) inhibitors^61^. Needless to say, the importance of dietary carbohydrate restriction vs increased dietary fat vs myocardial ketone metabolism in relation to heart failure treatment requires further study.

## Conclusions and Limitations of Study

In conclusion, we show that the MPC is deactivated in failing human and mouse hearts and that cardiac deletion of MPC2 in mice results in progressive cardiac hypertrophy and dilated heart failure. Interestingly, heart failure in CS-MPC2-/- mice could be prevented or even reversed by feeding a ketogenic or high fat diet and a ketogenic diet also limited remodeling in wild-type mice after TAC+MI surgery. Available data suggest that these improvements may be predominantly mediated by increasing myocardial FAO and limiting provision of carbohydrate, rather than enhancing ketone metabolism. However, we cannot definitively assess whether increasing ketosis alone improves cardiac function in CS-MPC2-/- mice, as it is difficult to achieve high levels of ketosis in the absence of increased dietary fat in rodents. Indeed, ketone body infusion was recently shown to improve cardiac function in dogs and humans with heart failure^29, 49^. Thus, some mechanistic aspects of the cause of heart failure observed in mice lacking MPC in the myocardium remain to be teased apart.

## METHODS

### Human Heart Tissue Collection

Human heart tissue was collected with written informed consent received from participants as part of an Institutional Review Board (IRB)-approved protocol (#201101858) at the Washington University School of Medicine. Failing human left ventricular heart tissue was obtained from the Washington University Translational Cardiovascular Tissue Core at the time of left ventricular assist device (LVAD) placement, or post-LVAD placement at the time of cardiac transplantation. Non-failing human heart tissue was obtained from Mid-America Transplant (St. Louis, MO) from hearts deemed unsuitable for transplantation due to donor age, non-occlusive coronary artery disease, or high-risk behavioral profile. The collected piece of cardiac tissue had fat removed, was rinsed in saline, and then was snap-frozen in liquid nitrogen and stored at -80°C until analyzed.

### Animals

All animal procedures were performed in accordance with National Institutes of Health guidelines and approved by the Institutional Animal Care and Use Committee at the Washington University School of Medicine. The use of mice conformed to guidelines set forth in the NIH’s *Guide for the Care and Use of Laboratory Animals* (National Academies Press, 2011).

Generation of mice with the *Mpc2* gene flanked by loxP sites has been described previously^24^. To create cardiac myocyte specific deletion, these mice were crossed with a knock-in mouse in which one allele of the myosin light chain 2v (Mlc2v) gene was replaced with Cre recombinase^62^, which was obtained from the Jackson Laboratory (Bar Harbor, ME). All mice were from a C57BL6/J background. Unless specifically noted, all experiments were performed with a mixture of male and female littermate mice. For the TAC+MI studies, 4-5 week-old wildtype C57BL6/J females were purchased from the Jackson Laboratory (Bar Harbor, ME).

### Animal Care

Mice were housed in a climate-controlled barrier facility maintained at 22-24°C and 40-60% humidity in ventilated cages with a 12-hour light/dark cycle with light period from 0600 to 1800 local time. *Ad-libitum* access to drinking water was provided by individual bottles in each cage. Mice were housed in cages with corn-cob bedding or switched to aspen chip bedding during special diet studies or fasting prior to sacrifice. All mice were group-housed, up to 5 mice per cage, with cloth nestlets to use for enrichment. Mice on special diets were also provided with a Nylabone (Central Garden & Pet Co., Neptune City, NJ) for both enrichment and to maintain teeth when fed soft, higher fat diets. With all diets, mice were provided *ad-libitum* access to food, except for a brief 4-hour fast prior to euthanasia. Unless specifically noted, all special diets were initiated at 6-weeks of age. All diets were provided on a wire rack above the cage bedding, with the exception of the ketogenic diet paste which was spread into a glass petri dish, placed on the bottom of the cage, and replaced every 2-3 days. Mice fed standard chow received PicoLab*®* Rodent Diet 20 (#5053, LabDiet, St. Louis, MO) which comprised of 62.1%kcal carbohydrate, 13.2%kcal fat, and 24.7%kcal protein. The refined low-fat (LF) diet was composed of 70%kcal carbohydrate, 10%kcal fat, and 20%kcal protein (D12450B, Research Diets, New Brunswick, NJ). Ketogenic diet (KD) was composed of 1.8%kcal carbohydrate, 93.4%kcal fat, and 4.7%kcal protein (F3666, Bio-Serv, Flemington, NJ). The medium-chain triglyceride (MCT) diet was composed of 37.9%kcal carbohydrate, 43%kcal fat (depleted of long-chain fatty acids), and 19.1%kcal protein (TD.00308, Envigo, Madison, WI). High-fat (HF) diet was composed of 20%kcal carbohydrate, 60%kcal fat, and 20%kcal protein (D12492, Research Diets, New Brunswick, NJ). Hearts were also analyzed from a cohort of WT mice fed a high trans-fat, fructose, cholesterol (HTF-C) diet (D09100301, Research Diets, New Brunswick, NJ) with or without insulin-sensitizing MPC-inhibitor MSDC-0602K treatment, which were previously published with respect to nonalcoholic steatohepatitis^26^. For the ketone ester (KE) diet experiment, control diet consisted of 63%kcal carbohydrate, 10%kcal fat, and 24%kcal protein (104403, Dyets, Bethlehem, PA), and the KE diet was composed of the same diet except 16.5%kcal of the carbohydrates were replaced with D-*β*-hydroxybutyrate-(R)-1,3 butanediol monoester “ketone ester” (16.5%kcal ketone-ester, 46.5%kcal carbohydrate, 10%kcal fat, and 24%kcal protein)(104404, Dyets, Bethlehem, PA). Control or KE diet were fed from 9-15 weeks of age.

Unless specifically noted, mice were euthanized after a 4 hour fast by CO2 asphyxiation and blood was collected via cannulation of the inferior vena cava into EDTA-treated tubes. Tissues were then excised, rinsed in PBS, weighed, and snap frozen in liquid nitrogen. Plasma was collected by spinning blood tubes at 8,000 x g for 8 minutes and then freezing the plasma supernatant in liquid nitrogen.

### Gene Expression Analysis

Levels of gene expression were determined by qPCR. Total RNA was extracted from snap frozen tissues using RNA-Bee (Tel-Test, Friendswood, TX). ∼50 mg of tissue was homogenized in RNA-Bee for 3-5 minutes using a 3 mm stainless steel bead at 30 Hz using a TissueLyser II (Qiagen, Hilden, Germany). RNA abundance and quality were assessed by Nanodrop (ThermoFisher Scientific, Waltham, MA). 1 μg of sample was then reverse transcribed into cDNA by Superscript VILO (ThermoFisher Scientific, Waltham, MA) using an Eppendorf Mastercycler*®* X50 thermocycler (Hauppauge, NY). Relative quantification of target gene expression was measured in duplicate using Power SYBR Green (ThermoFisher Scientific, Waltham, MA), using an ABI 7500 Real-Time PCR System (ThermoFisher Scientific, Waltham, MA). Target gene Ct values were normalized to reference gene (*Rplp0*) Ct values by the 2^-^*^ΔΔ^*^Ct^ method. Oligonucleotide primer sequences used for qPCR are listed in Supplemental Table 3.

### Western Blotting and Protein Expression Analysis

Protein extracts were prepared by homogenizing ∼50 mg of frozen tissue in an NP-40-based lysis buffer (15 mM NaCl, 25 mM Tris Base, 1 mM EDTA, 0.2% NP-40, 10% glycerol) supplemented with 1X cOmplete*™* protease inhibitor cocktail (Roche, Basel, Switzerland) and a phosphatase inhibitor cocktail (1 mM Na3VO4, 1 mM NaF, and 1mM PMSF). Tissue was homogenized in this buffer for 3-5 minutes using a pre-chilled 3 mm stainless steel bead at 30 Hz using a TissueLyser II (Qiagen, Hilden, Germany). Protein concentrations were measured using a Pierce MicroBCA kit (ThermoFisher Scientific, Waltham, MA), and detected with a BioTek Synergy plate reader and Gen5 software (BioTek Instruments, Winooski, VT). 50 μg of protein lysate was electrophoresed on precast Criterion 4-15% polyacrylamide gels (Biorad, Hercules, CA), and transferred onto 0.45 μm Immobilon PVDF membranes (MilliporeSigma, St. Louis, MO). Membranes were blocked with 5% Bovine Serum Albumin (Sigma, St. Louis, MO) in TBS-T for at least 1 hour.

Primary antibodies were then used at 1:1000 (or 1:5000 for VLCAD and LCAD) in 5%BSA-TBS-T overnight while rocking at 4°C. Antibodies for human MPC1 and MPC2 were from Cell Signaling (Danvers, MA), while anti-mouse MPC1 and MPC2 antibodies were a gift from Dr. Michael Wolfgang^24, 63, 64^. Antibodies for VLCAD^65^, LCAD^66^, and MCAD^67^ were gifts from Drs. Daniel Kelly or Arnold Strauss. Anti-CPT1B antibody was from Alpha Diagnostic

International (San Antonio, TX). Anti-BDH1 antibody was from ThermoFisher Scientific (Waltham, MA). Phospho-ERK1/2 (Thr202/Tyr204), Total ERK1/2, Phospho-AMPK*α* (Thr172), Total AMPK*α*, Phospho-mTOR (Ser2448), Total mTOR, Phospho-S6 Ribosomal Protein (Ser235/236), and Total S6 Ribosomal Protein were from Cell Signaling (Danvers, MA). Anti-*α*-Tubulin and *β*-Actin antibodies were from Sigma (St. Louis, MO). After primary antibody incubation, membranes were washed 3-5X for 5 min in TBS-T, and probed with IRDye secondary antibodies at 1:10,000 (Li-Cor Biosciences, Lincoln, NE) in 5%BSA-TBS-T for 1 hour. Membranes were imaged on an Odyssey*®* imaging system and analyzed with Image Studio*™* Lite software (Li-Cor Biosciences, Lincoln, NE). If needed for alignment, blot images were rotated with NIH ImageJ (Bethesda, MD).

### Mitochondrial Isolation and High Resolution Respirometry

Mitochondria were isolated by differential centrifugation from whole mouse hearts by homogenization with 10 passes of a glass-on-glass Dounce on ice with 4 mL of buffer containing 250 mM sucrose, 10 mM Tris Base, and 1 mM EDTA (pH 7.4). Homogenates were then spun at 1,000 x g for 5 min at 4°C to pellet nuclei and undisrupted cell debris. The supernatant was then spun at 10,000 x g for 10 min to pellet the mitochondrial fraction. The mitochondrial pellet was washed twice in homogenization buffer minus the EDTA with 10,000 x g 10 min spins. After the final wash, mitochondrial pellets were solubilized in ∼150 μL Mir05 respiration buffer (0.5 mM EGTA, 3 mM MgCl2, 60 mM Lactobionic acid, 20 mM Taurine, 10 mM KH2PO4, 20 mM HEPES, 110 mM sucrose, and 1 g/L fatty acid free Bovine Serum Albumin; pH 7.1). Mitochondrial protein concentration was then measured using a Pierce MicroBCA kit (ThermoFisher Scientific, Waltham, MA), and detected with a Synergy plate reader and Gen5 software (BioTek Instruments, Winooski, VT).

To measure oxygen consumption rates, 50 μg of mitochondrial protein was added to each 2 mL chamber of an Oxygraph O2k equipped with DatLab software (Oroboros Instruments, Innsbruck, Austria). Substrates used to assess pyruvate-stimulated respiration were 5 mM sodium pyruvate, 2 mM malate, 2.5 mM ADP+^Mg2+^, and then 10 μM UK-5099. To assess respiration on other substrates, 50 μM palmitoyl-DL-carnitine and 2 mM malate ± 2.5 mM ADP+^Mg2+^, or 10 mM glutamate and 2 mM malate ± 2.5 mM ADP+^Mg2+^, or 5 mM succinate + 2.5 mM ADP+^Mg2+^ were used. Oxygen consumption rates were measured as pmol O2/s/mg mitochondrial protein.

### Blood and Plasma Metabolite and Hormone Measurements

Immediately prior to euthanasia, a snip of the tail was made with a razor blade and a drop of mixed venous blood was used to measure blood glucose using a Contour Next EZ (Bayer Ascensia Diabetes Care, Parsippany, NJ) glucometer. A second drop of blood was then used to measure blood lactate concentrations using a Lactate Plus meter (Nova Biomedical, Waltham, MA). Plasma insulin concentrations were measured from 10 μL of plasma by Singulex Erenna*®* assay (Sigma, St. Louis, MO) performed by the Washington University Core Lab for Clinical Studies. Total ketone bodies were measured from 4 μL plasma using the Total Ketone AutoKit (FujiFilm Wako, Mountain View, CA) according to kit directions. Optic density (OD) at 405 and 600 nm were measured every minute for 5 minutes, and absorbance changes were normalized to a 300 μM standard.

Free fatty acids were measured from 2 μL plasma using a non-esterified fatty acid kit according to manufacturer’s directions (FujiFilm Wako, Mountain View, CA). OD at 560 nm was measured and normalized to a standard curve. Triglycerides were measured from 5 μL and cholesterol was measured from 2 μL plasma using Infinity assay kits according to manufacturer’s directions (ThermoFisher Scientific, Waltham, MA). OD and 540 was measured and related to the OD of a standard curve. Free glycerol was measured from 10 μL plasma and measured according to manufacturer’s directions (Sigma, St. Louis, MO) for OD at 540 nm. OD for all assays was measured in clear 96-well plates using a Synergy plate reader and Gen5 software (BioTek Instruments, Winooski, VT).

### Targeted Metabolomics for Amino Acids, Acylcarnitines, Organic acids, and Short Chain Acyl- CoAs

Mice used for targeted metabolomic analyses were fasted for 3 hours, anesthetized with 100 μg/g sodium pentobarbital injected i.p., and euthanized by excision of the beating heart.

Hearts were snap frozen in liquid nitrogen and stored at -80°C until they were collectively processed and analyzed. Flash frozen hearts were pulverized to a fine powder in a liquid nitrogen chilled percussion mortar and pestle and weighed into pre-chilled 2ml tubes. A chilled 5mm homogenizing bead was added to samples and tissue was diluted to 50 mg/ml with 50% acetonitrile containing 0.3% formate (for acylcarnitines, amino acids, and organic acids) or isopropanol/phosphate buffer (for CoAs), homogenized for 2 min at 30 Hz using a TissueLyser II (Qiagen, Hilden, Germany) and aliquoted for metabolite assays. For all metabolite analyses, tissues and homogenates were kept on ice, centrifuged at 4°C, and when ready to measure, were placed in an autosampler kept at 4°C.

Amino acids and acylcarnitines were analyzed by flow injection electrospray ionization tandem mass spectrometry and quantified by isotope or pseudo-isotope dilution similar to previous^68–70^, which are based on methods developed for fast ion bombardment tandem mass spectrometry^71^. Extracted heart samples were spiked with a cocktail of heavy-isotope internal standards (Cambridge Isotope Laboratories, Tewksbury, MA; or CDN Isotopes, Pointe-Claire, Canada) and deproteinated with methanol. The methanol supernatants were dried and esterified with either acidified methanol or butanol for acylcarnitine or amino acid analysis, respectively. Mass spectra for acylcarnitine and amino acid esters were obtained using precursor ion and neutral loss scanning methods, respectively. The spectra were acquired in a multi-channel analyzer (MCA) mode to improve signal-to-noise. The data were generated using a Waters TQ (triple quadrupole) detector equipped with Acquity^TM^ UPLC system and a data system controlled by MassLynx 4.1 operating system (Waters, Milford, MA). For the amino acids analysis, the mass spectrometer settings were as follows: ionization mode - positive electrospray, capillary voltage - 3.6 V, cone voltage - 14 V, extractor voltage - 2 V, RF lens voltage - 0.1 V, collision energy - 14-25 V, source temperature - 130℃, desolvation temperature - 200℃, desolvation gas flow - 550 L/hr, and cone gas flow - 50 L/hr. For the acylcarnitine analysis, the mass spectrometer settings were as follows: ionization mode - positive electrospray, capillary voltage - 3.5 V, cone voltage - 25 V, extractor voltage - 2 V, RF lens voltage - 0.1 V, collision energy - 30 V, source temperature - 130℃, desolvation temperature - 200℃, desolvation gas flow - 550 L/hr, and cone gas flow - 50 L/hr. Ion ratios of analyte to respective internal standard computed from centroided spectra are converted to concentrations using calibrators constructed from authentic aliphatic acylcarnitines and amino acids (Sigma, St. Louis, MO; Larodan, Solna, Sweden) and Dialyzed Fetal Bovine Serum (Sigma, St. Louis, MO).

Organic acids were analyzed by capillary gas chromatography/mass spectrometry (GC/MS) using isotope dilution techniques employing Trace Ultra GC coupled to ISQ MS operating under Xcalibur 2.2 (ThermoFisher Scientific, Austin, TX)^72^. The mass spectrometer settings were as follows: ionization mode - electron ionization, ion source temperature - 250℃, and the transfer line temperature - 275℃. The supernatants of tissue homogenates were spiked with a mixture of heavy isotope labeled internal standards and the keto acids were stabilized by ethoximation. The organic acids were acidified and extracted into ethyl acetate. The extracts were dried and derivatized with N,O-bis(Trimethylsilyl) trifluoroacetamide. The organic acids were quantified using ion ratios determined from single ion recordings of fragment ions which are specific for a given analyte and its internal standard. These ratios were converted to concentrations using calibrators constructed from authentic organic acids (Sigma, St. Louis, MO). The heatmap for acylcarnitines was generated by shinyheatmap^73^.

Short chain acyl CoA were analyzed by LC-MS/MS using a method based on a previously published report^74^. The extracts were spiked with ^13^C2-Acetyl-CoA, centrifuged and filtered through the Millipore Ultrafree-MC 0.1 µm centrifugal filters before being injected onto the Chromolith FastGradient RP-18e HPLC column, 50 x 2 mm (MilliporeSigma, St. Louis, MO) and analyzed on a Waters Xevo TQ-S triple quadrupole mass spectrometer coupled to an Acquity UPLC system (Waters, Milford, MA). The mass spectrometer settings were as follows: ionization mode - positive electrospray, capillary voltage - 3.7 V, cone voltage - 50 V, source offset voltage - 50 V, collision energy - 28 V, dwell time - 0.06 seconds, desolvation temperature - 500℃, desolvation gas flow - 600 L/hr, cone gas flow - 150 L/hr, and nebulizer pressure - 7 bar. The following MRM transitions were monitored: Acetyl- CoA - 810.2->303.1, ^13^C2-Acetyl-CoA – 812.2->305.1, Succinyl-CoA - 868.2 ->361.1, and Malonyl-CoA – 854.2->347.1.

### Mouse Echocardiography

*In vivo* cardiac size and function were measured by echocardiography performed with a Vevo 2100 Ultrasound System equipped with a 30-MHz linear-array transducer (VisualSonics Inc, Toronto, Ontario, Canada)^32^. Mice were lightly anesthetized by i.p. injection of 0.005 ml/g of 2% Avertin (2,2,2-tribromoethanol; Sigma, St. Louis, MO). If required, one-fifth of the initial dose was given as a maintenance dose at regular intervals. Hair was removed from the left anterior chest by shaving, and mice were then placed onto a warming pad in a left lateral decubitus position. Normothermia (37°C) was maintained and monitored by a rectal thermometer. Ultrasound gel was applied to the chest and care was taken to maintain adequate transducer contact while avoiding excessive pressure on the chest. Two-dimensional and M-mode images were obtained in the long- and short-axis views. Images were retrieved off- line and analyzed using the Vevo LAB software package (VisualSonics Inc, Toronto, Ontario, Canada). Measurements were averaged from three separate images for each mouse. LV volumes were calculated from M-mode measurements using standard techniques^75, 76^. Immediately after completion of imaging, mice were allowed to recover from anesthesia on a warming pad and returned to their home cage. For echocardiography during the ketone ester diet experiment, procedures were the same as above except mice were anesthetized by 1-2% inhaled isoflurane, and imaging was performed with a Vevo 770 Ultrasound System equipped with a 30-MHz linear-array transducer (VisualSonics Inc, Toronto, Ontario, Canada).

### Histology

Short-axis slices of the LV were fixed in 10% neutral buffered formalin overnight and processed by the Anatomic and Molecular Pathology core laboratory of Washington University. The short-axis heart slices were embedded in paraffin blocks and sectioned onto glass slides. Slides were then stained for either Hematoxylin and Eosin (H&E) or Mason’s trichrome stains.

### Body Composition Analysis

Mouse body composition analysis was performed using an EchoMRI 3-in-1 system (EchoMRI*™*, Houston, TX). Briefly, after machine calibration with an olive oil standard, conscious mice were restrained in a plastic tube and placed into the instrument bore. Fat, lean, free water, and total water masses were then determined. Imaging required <5 min per mouse, and following imaging, mice were immediately placed back into their home cage.

### TAC+MI Surgically-Induced Heart Failure Model

7-week old female WT C57BL/6J mice (Jackson Laboratory, Bar Harbor, ME) were subjected to TAC+MI surgery as performed previously^32, 77^. Mice were anesthetized with 100 mg/kg ketamine and 10 mg/kg xylazine injected i.p. and were then restrained supine, intubated, and ventilated with a respirator (Harvard Apparatus, Holliston, MA). After shaving of the left anterior chest, the intercostal muscles were dissected, and aorta identified and freed by blunt dissection. 7-0 silk suture was placed around the transverse aorta and tied around a blunt 26-gauge needle. The needle was then removed after placement of the constrictor. Immediately following the first procedure, the LV and left main coronary artery system were exposed and the apical portion of the LAD was ligated with 9-0 silk suture. The surgical incision was closed, and the mice were recovered on a warmer until arousal from anesthesia whence they were returned to their home cage. All surgeries were performed in under 20 minutes.

Two weeks post TAC+MI surgery, mice were imaged by echocardiography with procedures similar to above and with modifications as performed previously^32^.

Echocardiography procedures specific to TAC+MI studies included Doppler ultrasound examination to measure the aortic flow gradient across the constriction^78^. With 2-D imaging guidance, the pulse wave Doppler sample volume was placed at the site of constriction. Aortic flow velocity was also measured proximal to the constriction near the aortic root to account for decreased cardiac output. Velocity time integral, mean, and peak gradients were measured, and the ratio of distal/proximal integral was calculated as an index of the aortic constriction gradient. Extent of the LV infarct region was also assessed by 2-D short-axis movies and measurement of akinetic segments as performed previously^32^. LV volumes were measured from parasternal long-axis views of the LV by the disk summation method^79^. After completion of the 2-week post TAC+MI echocardiography, images were analyzed, and mice were assigned into LF- or KD-fed groups so that both groups started with roughly equal cardiac remodeling and dysfunction. After consuming LF or KD diets for 2 weeks (4-weeks total post TAC+MI), echocardiography was repeated and mice were euthanized the following day by CO2 asphyxiation after a 4 hour fast for collection of plasma and tissues.

### Ketone Body Injection

12-week old CS-MPC2-/- mice underwent echocardiography as detailed above and were then randomized into two groups to receive daily i.p. injections of either saline vehicle or 10 mmol/kg R-3-hydroxybutyric acid sodium salt (*β*HB) (Sigma, St. Louis, MO), which was pH’d to ∼7.0. After 2 weeks of daily i.p. injection, echocardiography was repeated following the same procedures as detailed above. The following day, mice received a final saline or *β*HB injection, were fasted for 4 hours, and were euthanized by CO2 asphyxiation for collection of plasma and tissues.

### Statistics

All data are presented as dot plots with mean +/- s.e.m., or as PRE-POST data points. Multiple comparisons were analyzed using a 2-way ANOVA with Tukey’s multiple-comparisons test. An unpaired, 2-tailed Student’s *t* test was used for comparison of 2 groups. A *P* value of less than 0.05 was considered statistically significant. Statistical analysis was performed using GraphPad Prism 8 software.

## ACKNOWLEDGEMENTS

Sadly, Dr. Richard (Bud) Veech passed away at the age of 84 during the preparation of this manuscript. We thank him for providing the ketone ester diet and his enthusiasm towards this project. This work was supported by core resources of the Nutrition Obesity Research Center (NORC) (P30 DK56341), Diabetes Research Center (DRC) (P30 DK020579), and Institute for Clinical and Translational Sciences (ICTS) (UL1 TR002345) at the Washington University School of Medicine. NIH grants K99/R00 HL136658 (to KSM), R01 HL133178 (to RWG), and R01 HL119225 and R01 DK104735 (to BNF) supported these studies.

## AUTHOR CONTRIBUTIONS

Conceptualization, KSM and BNF; Methodology, KSM, AK, CJW, TMS, TRK, ORI, DMM, and BNF; Investigation, KSM, AK, CJW, TRK, ORI, DRK, and KDP; Resources, MTK, RLV, BJD, and RWG; Writing – original draft, KSM and BNF; Writing – review & editing, KSM, AK, CJW, TMS, TRK, ORI, DMM, DRK, KDP, MTK, BJD, RWG, and BNF; Funding Acquisition, KSM and BNF; Supervision, KSM, AK, CJW, and BNF.

