## Supplemental Figures 1-8 for "Ketogenic Diet Prevents Heart Failure from Defective Mitochondrial Pyruvate Metabolism"

Supplemental Fig. 1

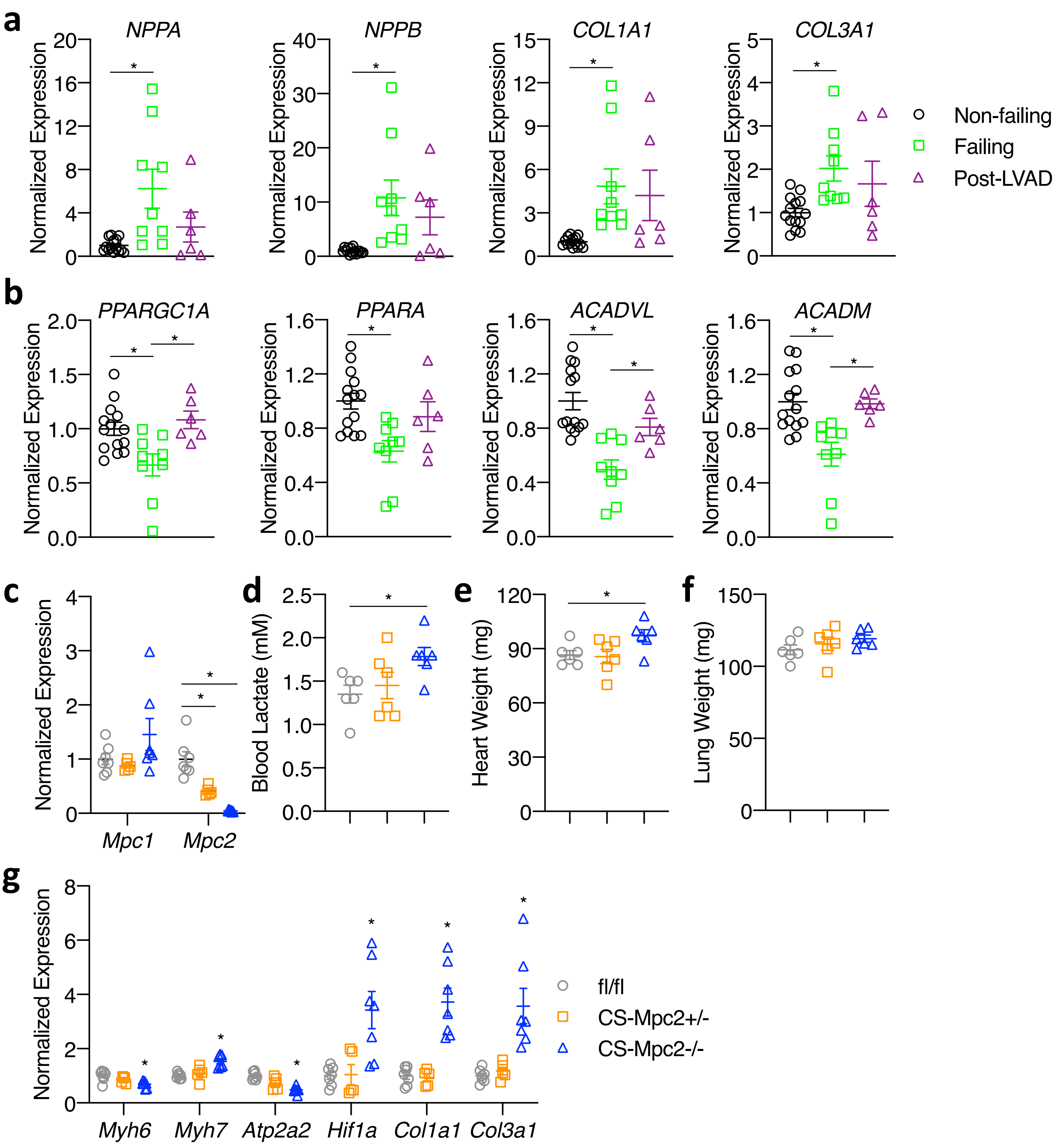


**Supplemental Fig. 1 – Related to Fig. 1. Human heart failure gene expression and characterization of 6-week old CS-MPC2-/- mice. a-b,** Gene expression from human non-failing, failing, and failing hearts after left ventricular assist device (LVAD) placement (n=6-14). **c,** Mouse heart gene expression for *Mpc1* and *Mpc2* (n=5-7). **d,** Blood lactate measured after a 4 h fast prior to sacrifice in 6-week old mice (n=6). **e-f,** Heart weight and lung weight normalized to body weight of 6-week old mice (n=6). **g,** Mouse heart gene expression of heart failure, hypertrophy, and fibrosis genes (n=6). Data are presented as mean ± s.e.m. within dot plot. Each data point represents one individual mouse or sample. Two-tailed unpaired Student's *t* test. $^{*}P < 0.05$, $^{**}P < 0.01$, $^{***}P < 0.001$.

### Supplemental Fig. 2

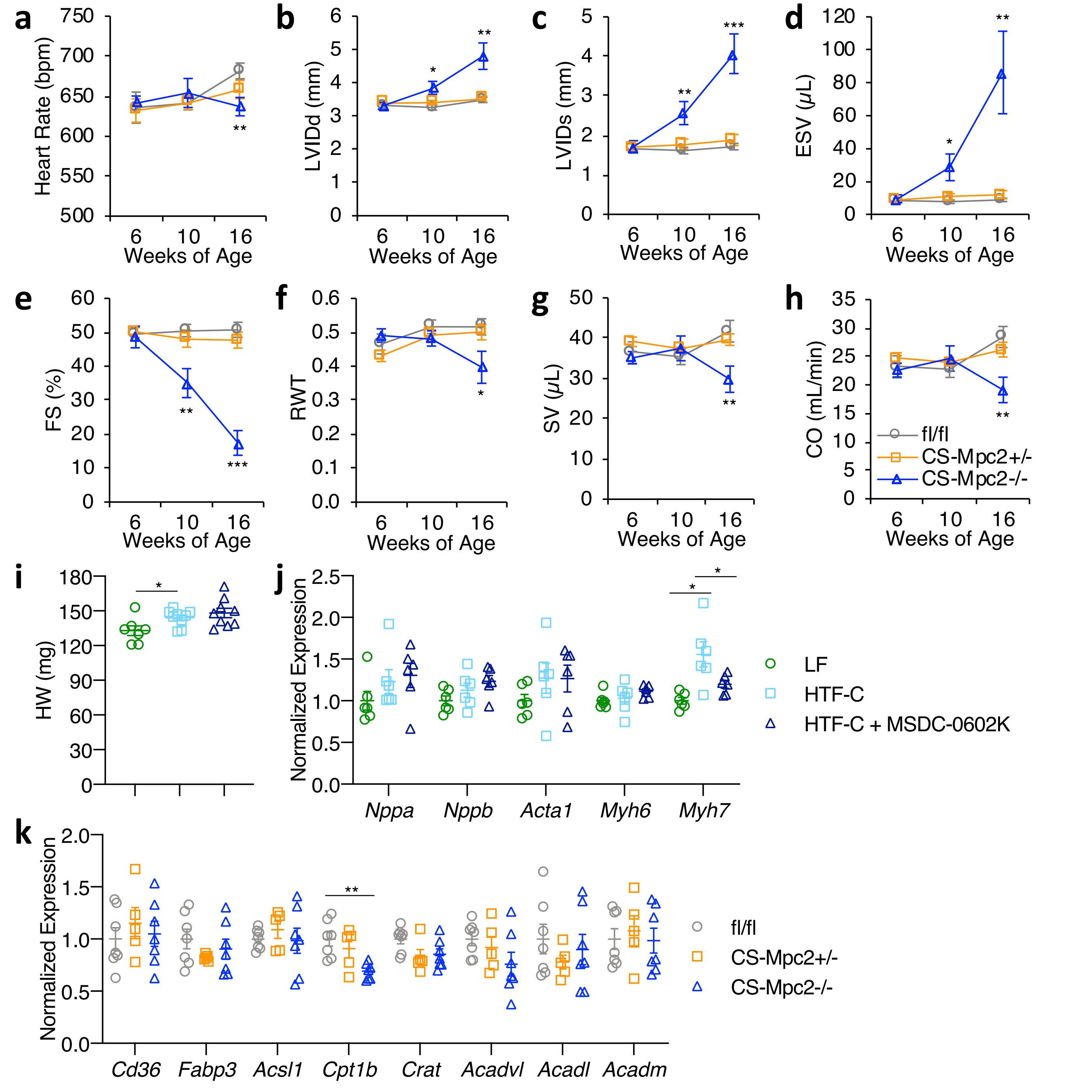


**Supplemental Fig. 2 – Related to Fig. 2. Heart failure develops in CS-MPC2-/- mice, but not CS-MPC2+/- or mice treated with the MPC inhibitor MSDC-0602K. a-h,** Serial echocardiography data of chow-fed mice at 6, 10, and 16 weeks of age. Left ventricular internal diameter at end diastole (LVIDd) and end systole (LVIDs), end systolic volume (ESV), fractional shortening (FS), relative wall thickness (RWT), stroke volume (SV), and cardiac output (CO). **i,** Heart weights from WT mice fed low fat (LF) diet or a high trans-fat, fructose, cholesterol (HTF-C) diet +/- 330 ppm MSDC-0602, an insulin-sensitizing MPC inhibitor (n=7-9). **j,** Heart gene expression of hypertrophy gene markers from WT mice fed LF, HTF-C, or HTF-C + MSDC-0602 diets (n=6). **k,** Heart gene expression for fatty acid transport and oxidation genes from 16-week old mice (n=6). Data are presented as mean ± s.e.m., or mean ± s.e.m. within dot plot. Each data point represents one individual mouse or sample. Two-tailed unpaired Student's *t* test. *$P < 0.05$, **$P < 0.01$, ***$P < 0.001$.

### Supplemental Fig. 3

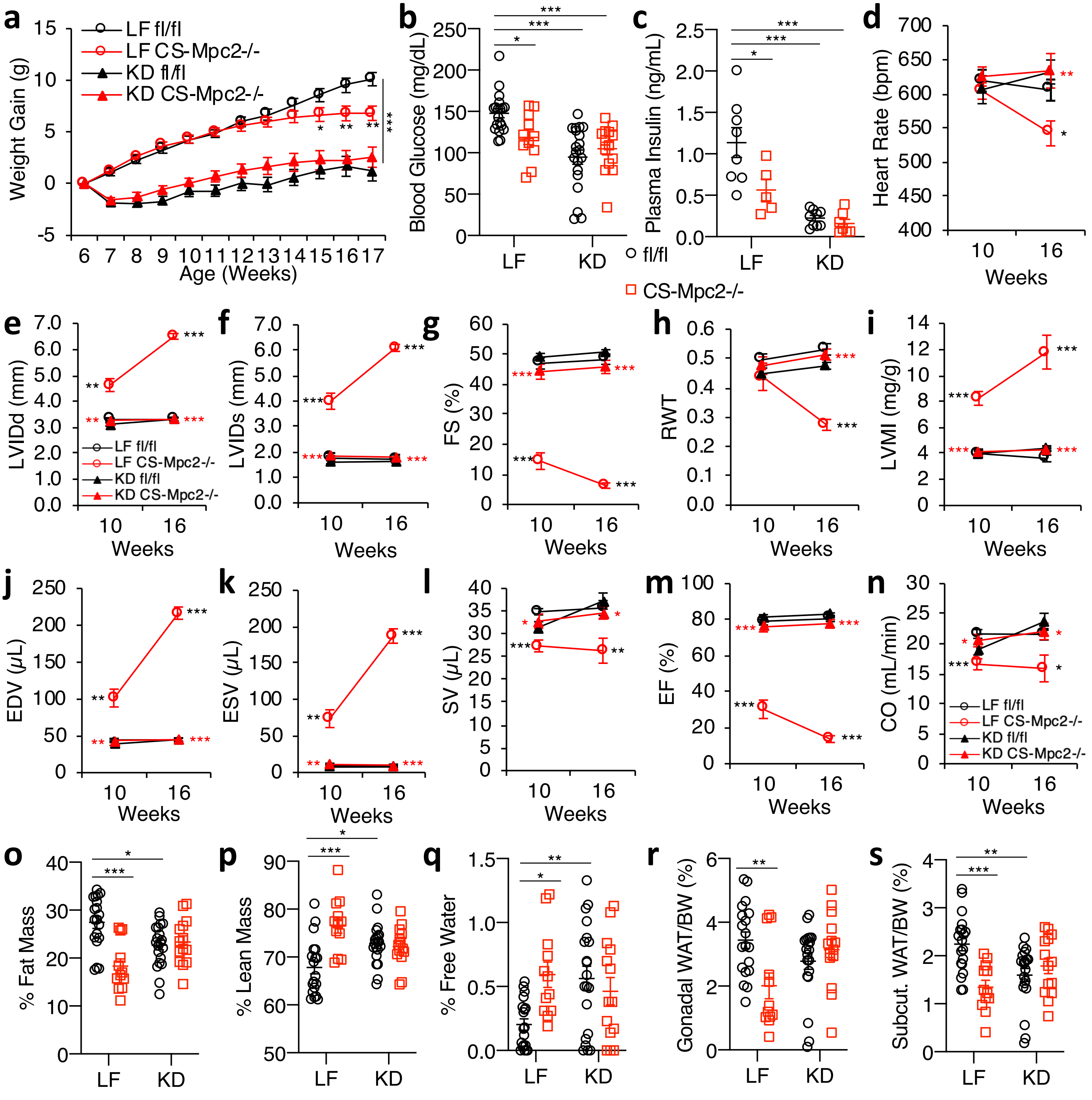


**Supplemental Fig. 3 – Related to Fig. 3. Ketogenic diet prevents heart failure in CS-MPC2-/- mice. a,** Body weight gain for mice fed low fat (LF) or ketogenic diet (KD) from 6-17 weeks of age (initial n=14-20). **b-c,** Blood glucose and plasma insulin measured after a 4 h fast immediately before sacrifice (n=7-11). **d-n,** Echocardiography data at 10 and 16 weeks of age. Left ventricular internal diameter at end diastole (LVIDd) and end systole (LVIDs), fractional shortening (FS), relative wall thickness (RWT), end diastolic volume (EDV), end systolic volume (ESV), stroke volume (SV), ejection fraction (EF), and cardiac output (CO) (n=7-11). **o-q,** % Fat mass, % lean mass, and % free water body composition measured by echoMRI (n=11-20). **r-s,** Gonadal and inguinal white adipose tissue (WAT) weights normalized to body weight (n=11-20). Data are presented as mean ± s.e.m. or mean ± s.e.m. within dot plot. Each data point in dot plot represents one individual mouse sample. Two-way ANOVA with Tukey's multiple comparisons test. $^{*}P < 0.05$, $^{**}P < 0.01$, $^{***}P < 0.001$. For **d-n**, black asterisks indicate between LF-fed fl/fl and CS-Mpc2-/-, red asterisks indicate between LF and KD (for CS-Mpc2-/-) for each echocardiography date.

Supplemental Fig. 4

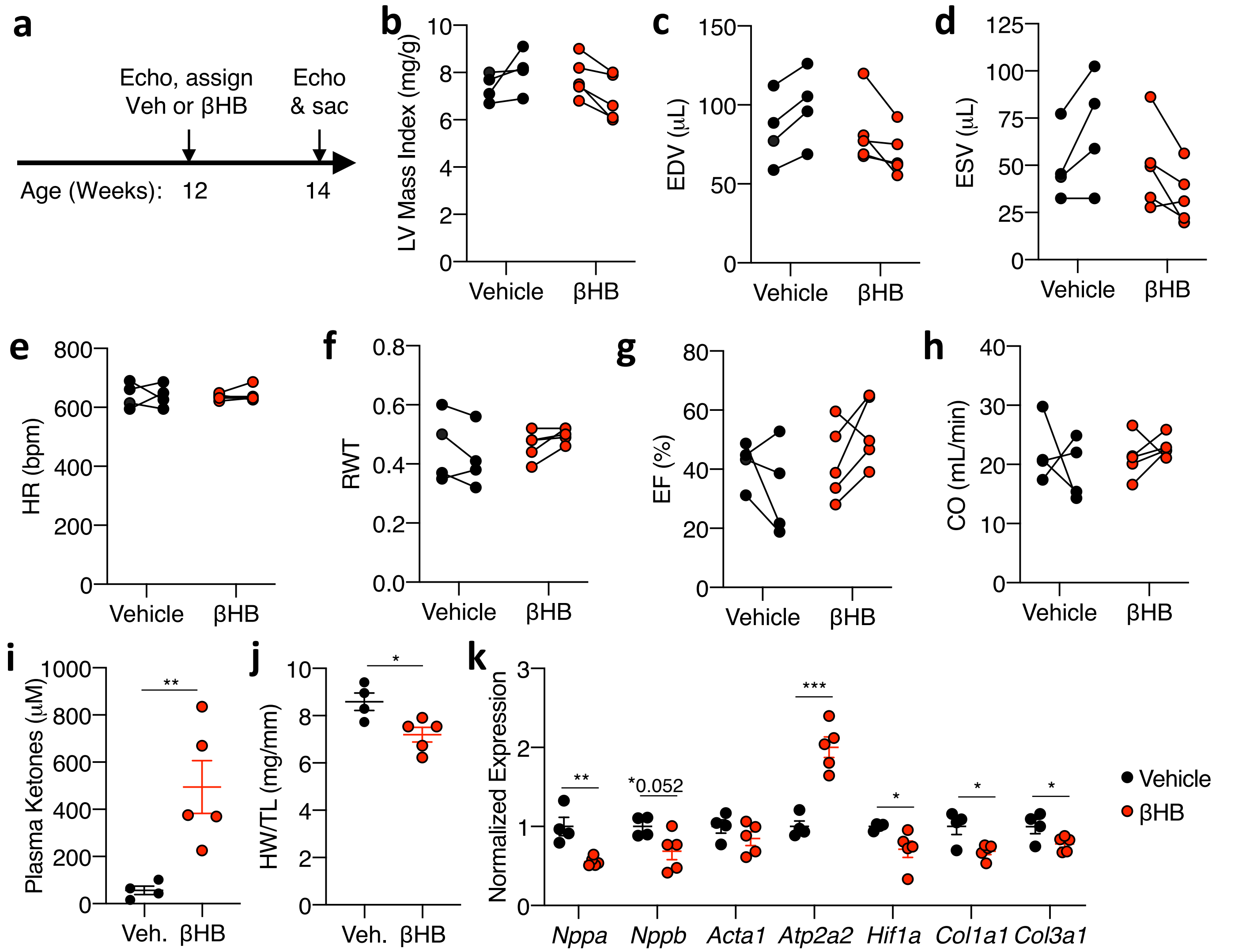


**Supplemental Fig. 4. Ketone body injection modestly reduces cardiac remodeling in CS-MPC2-/- mice. a,** Timeline for β-hydroxybutyrate (βHB) injection experiment in which CS-MPC2-/- mice were injected i.p. with saline vehicle or 10 mmol/kg βHB daily from 12 to 14 weeks of age. **b-h,** Echocardiography measurements before and after 2 weeks of daily i.p. injection of saline vehicle (Veh) or 10mmol/kg β-hydroxybutyrate. Left ventricular (LV) mass index, end-diastolic volume (EDV), end-systolic volume (ESV), heart rate (HR), relative wall thickness (RWT), ejection fraction (EF), and cardiac output (CO) (n=4-5). **i,** Plasma total ketone body concentrations (n=4-5). **j,** Heart weight normalized to tibia length (n=4-5). **k,** Gene expression markers of hypertrophy, heart failure, and fibrosis from hearts after 2 weeks of daily vehicle or βHB treatment (n=4-5). Data presented either as PRE-POST, or mean ± s.e.m. shown within dot plot. Each symbol represents an individual sample. Two-tailed unpaired Student's *t* test. *$P < 0.05$, **$P < 0.01$, ***$P < 0.001$.

Supplemental Fig. 5

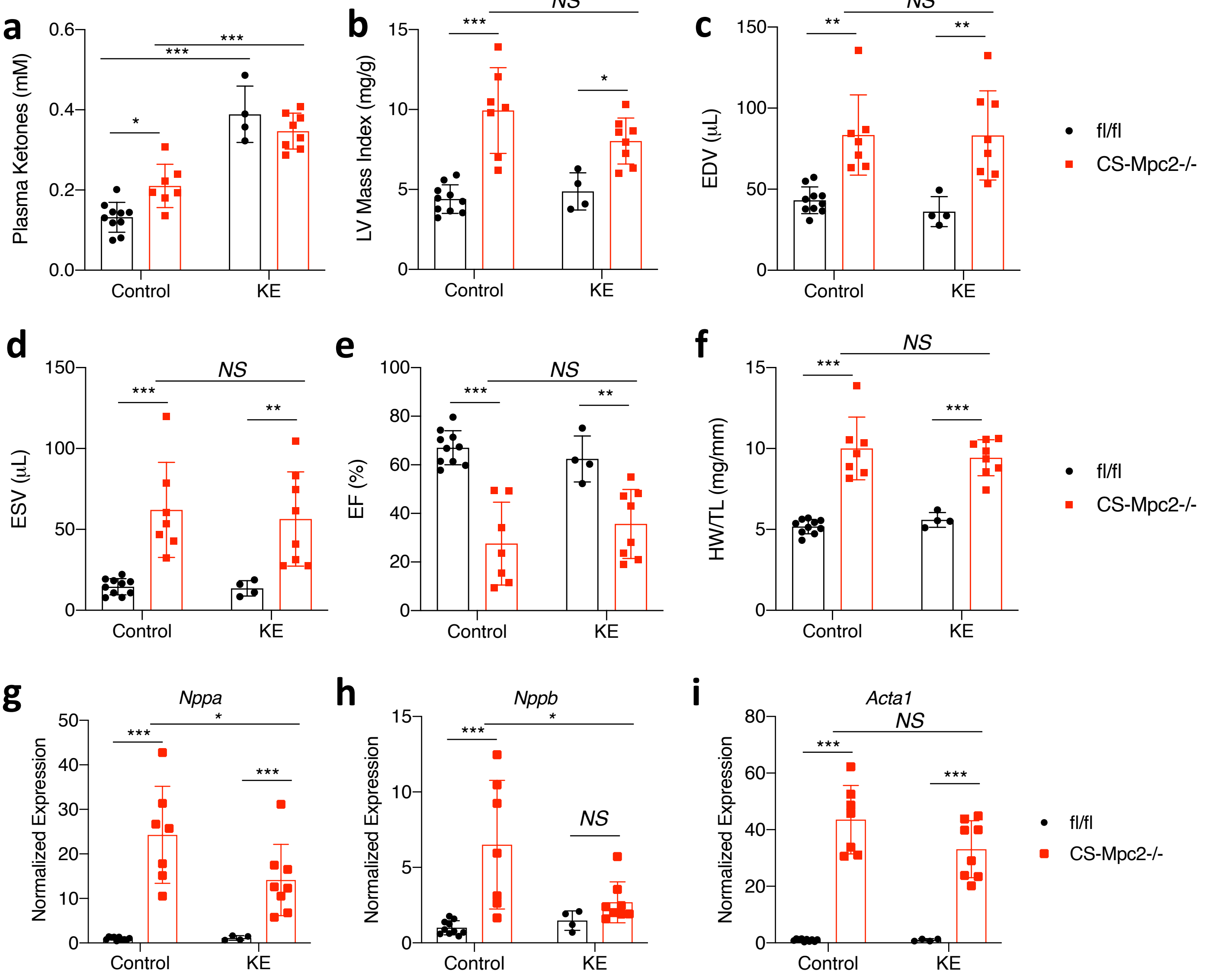


**Supplemental Fig. 5. Ketone ester diet does not improve cardiac remodeling or function in CS-MPC2-/- mice. a,** Plasma ketone bodies measured from mice fed either control or ketone ester (KE)-supplemented diet (n=4-10). **b-e,** Echocardiography measurements after 6 weeks of KE diet feeding. Left ventricular (LV) mass index, end-diastolic volume (EDV), end-systolic volume (ESV), and ejection fraction (EF)(n=4-10). **f,** Heart weight normalized to tibia length (n=4-10). **g-i,** Cardiac gene expression markers of hypertrophy and heart failure (*Nppa*, *Nppb*, *Acta1*)(n=4-10). Data presented as mean ± s.e.m. shown within dot plot. Each symbol represents an individual sample. Two-way ANOVA with Tukey's multiple comparisons test. *$P < 0.05$, **$P < 0.01$, ***$P < 0.001$.

Supplemental Fig. 6

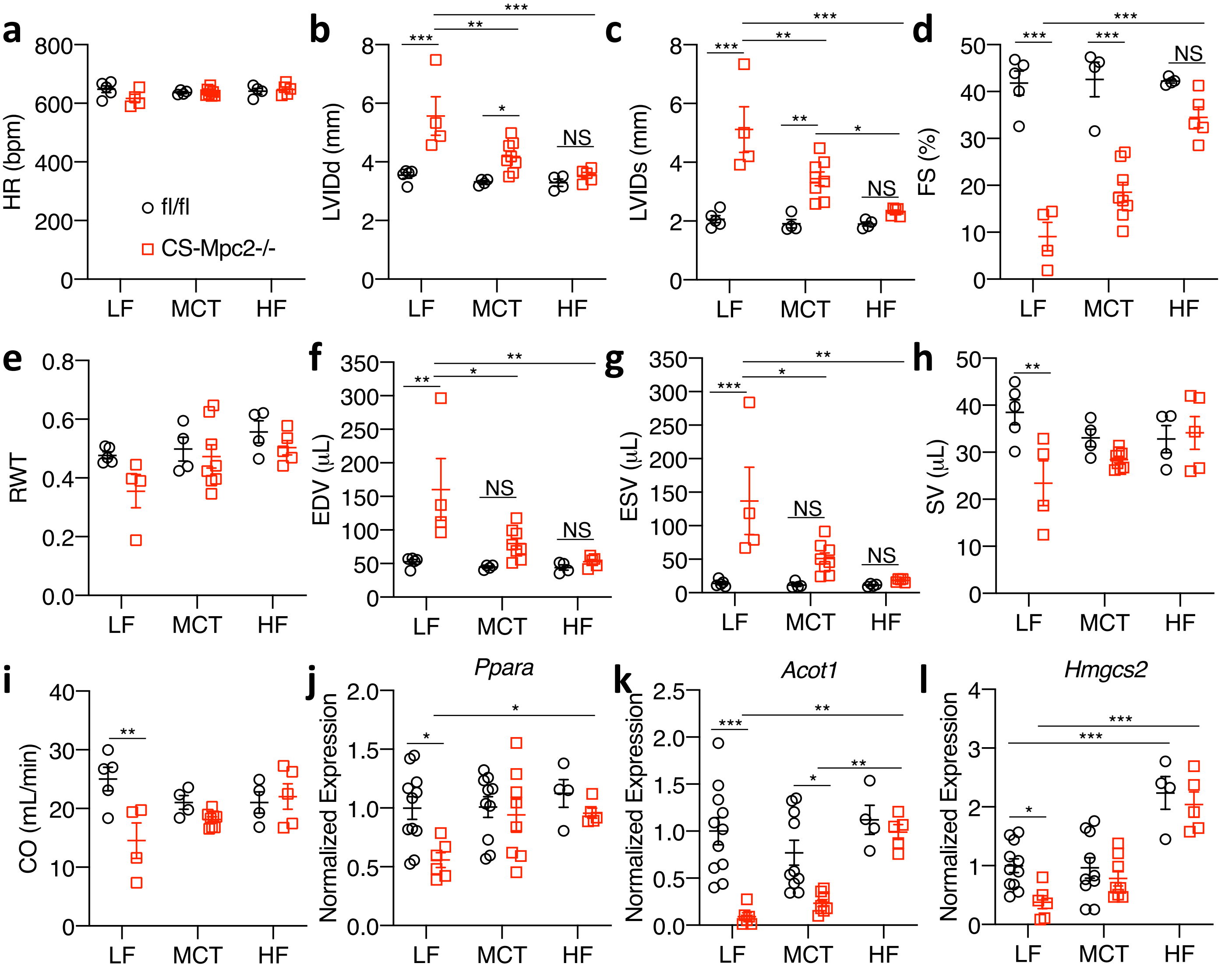


**Supplemental Fig. 6 – Related to Fig. 6. High fat diets also greatly improve cardiac remodeling and function of CS-MPC2-/- mice. a-I,** Echocardiography measurements taken at 16 weeks of age after 10 weeks of low fat (LF), medium chain triglyceride (MCT), or high-fat (HF) feeding. Left ventricular internal diameter at end diastole (LVIDd) and end systole (LVIDs), fractional shortening (FS), relative wall thickness (RWT), end diastolic volume (EDV), end systolic volume (ESV), stroke volume (SV), and cardiac output (CO) (n=4-11). **j-l,** Gene expression for *Ppara* and it's targets *Acot1* and *Hmgcs2* from mouse hearts (n=4-11). Data are presented as mean ± s.e.m. within dot plot. Each data point represents an individual mouse. Two-way ANOVA with Tukey's multiple comparisons test. $^{*}P < 0.05$, $^{**}P < 0.01$, $^{***}P <$ 0.001.

### Supplemental Fig. 7

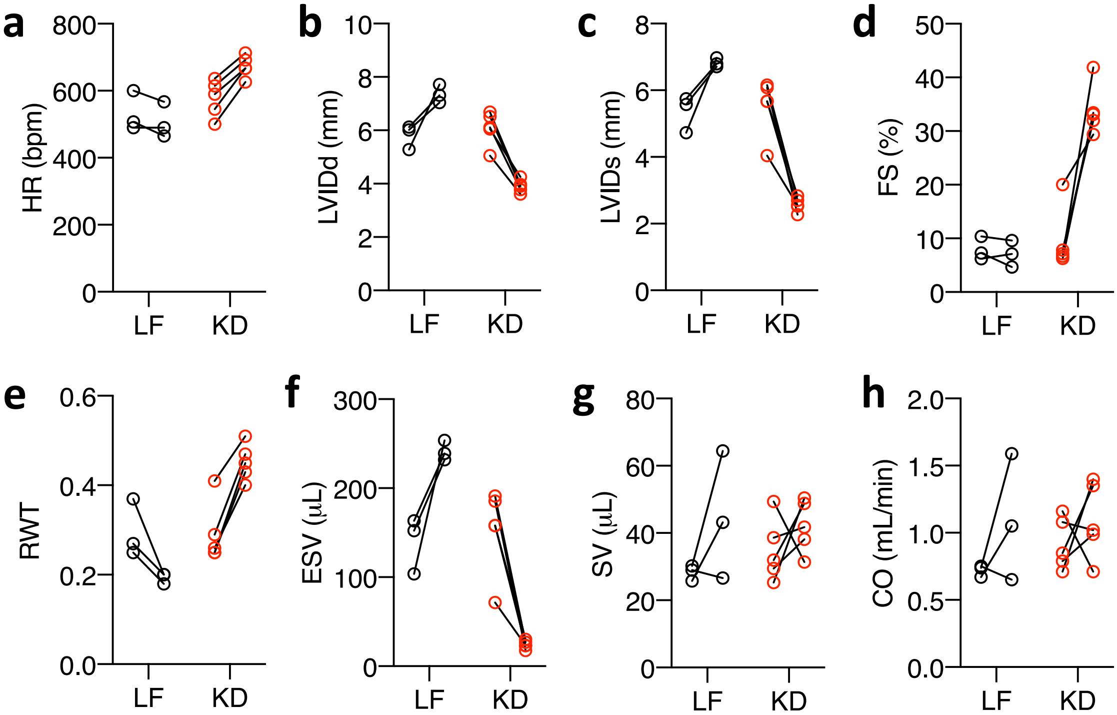


**Supplemental Fig. 7 – Related to Fig. 7. Ketogenic diet reverses heart failure in CS-MPC2-/- mice. a-h,** Echocardiography measurements before and after 3 weeks of LF or KD-feeding in 16 week old CS-MPC2-/- mice with established heart failure (n=3-5). Data are presented as PRE-POST. Each data point represents an individual mouse.

### Supplemental Fig. 8

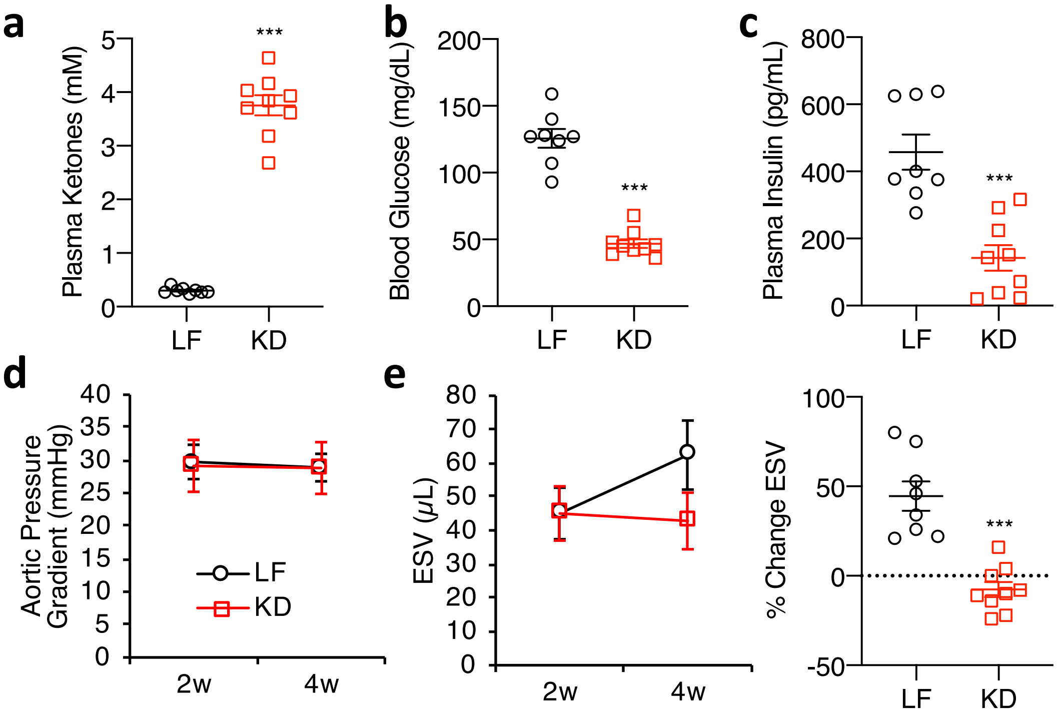


**Supplemental Fig. 8 – Related to Fig. 8. Ketogenic diet prevents cardiac remodeling in TAC+MI model of heart failure. a-c,** Plasma ketones, blood glucose, and plasma insulin measured after 2 weeks of LF or KD feeding (n=8-9). **d-e,** Echocardiography measures of mean aortic pressure gradient and end systolic volume (ESV) at 2 and 4 weeks post TAC+MI surgery (n=8-9). Data are presented as mean ± s.e.m. within dot plot. Each symbol on dot plot represents an individual sample. Two-tailed unpaired Student's *t* test. *$P < 0.05$, **$P < 0.01$, ***$P < 0.001$.
